# Chimerophore antibiotics: engineered multimodal host defense peptides

**DOI:** 10.64898/2026.08.06.743220

**Authors:** Liz Armas-Egas, Sandra Lagler, Sven Panke, Martin Held

## Abstract

Ribosomally synthesized host defense peptides (HDPs) are promising candidates for novel antibiotics. However, non-lytic HDPs, which target intracellular machinery, remain underexploited due to limitations including low potency in serum and narrow activity spectra. To enhance their therapeutic profile, we fused non-lytic HDPs generating “chimerophores” with multimodal mechanisms of action (MOAs). Using a high-throughput self-screening platform (Me^x^), we synthesized and evaluated a combinatorial library of 99,235 variants, identifying over 30,300 active chimerophores, vastly expanding the functional space of chimeric HDPs. Functional screening of 18 chimerophores revealed candidates with potent, broad-spectrum activity displaying serum-tolerance, low cytotoxicity, orthogonal uptake pathways and multimodality, such as simultaneously targeting of ribosomes and DNA. Integrating these distinct mechanisms into a single molecule allowed lead candidate cp9 to suppress the emergence of resistance in *Pseudomonas aeruginosa*, establishing a scalable platform for the systematic engineering of next-generation multimodal antibiotics.

## Introduction

The stagnant antibiotic pipeline cannot keep pace with rising antimicrobial resistance, rendering bacterial infections a resurgent global threat. Ribosomally synthesized host defense peptides (HDPs) constitute an evolutionarily conserved class of molecules for innate defense across multicellular organisms. Among other functions, HDPs efficiently combat microbial infections either by directly disrupting the membranes of invading pathogens (lytic) or by inhibiting intracellular processes essential for pathogen survival (non-lytic)^1^. Because HDPs bind to large molecular surfaces of highly conserved targets (e.g., the bacterial membrane or 70S ribosomes) for which a single point mutation may be insufficient to abolish peptide binding, they possess an inherently higher barrier to resistance emergence via target modification than single-enzyme inhibiting antibiotics ^2–4^ . Due to this high barrier to resistance and their diverse mechanisms of action (MOAs), HDPs have emerged as a promising reservoir for the development of new antibiotic classes. However, they remain an essentially unexploited resource, as their transition into clinical development as systemic drugs has been hampered by critical obstacles, such as off-target toxicity and low bioavailability^5^.

Off-target toxicity frequently stems from the same physicochemical properties that confer their potent antimicrobial activities: high cationicity and amphipathicity^6,7^. These features drive antimicrobial activity via self-promoted uptake, initiated by electrostatic interactions with the negatively charged, solvent-exposed bacterial cell envelope components (e.g., lipopolysaccharides (LPS) or lipoteichoic acids (LTA)) followed by peptide insertion into the hydrophobic core of the phospholipid bilayer^8–10^. Because the outer leaflet of mammalian lipid bilayers is zwitterionic and enriched with neutral cholesterol, many cationic HDPs preferentially target bacterial cells. Nevertheless, the structural similarities between mammalian and bacterial membranes can be sufficient to allow spontaneous partitioning into host membranes at bactericidal concentrations, resulting in hemolysis or host-cell damage^8,9^. This introduces a critical therapeutic dilemma for lytic HDPs as the peptide saturation concentrations required to achieve bacterial membrane disruption are frequently high enough to trigger host cell membrane destabilization^5,11^.

In contrast, non-lytic cationic HDPs possess lower amphipathicity and translocate across bacterial lipid bilayers without disrupting them. Once inside the cytoplasm, they selectively bind macromolecular targets such as bacterial ribosomes, DNA, or protein-folding chaperones, thereby inhibiting essential pathways and processes. Because their intracellular targets are structurally distinct from mammalian orthologs, and because no membrane-disruptive critical concentration is required for achieving an antibacterial effect, non-lytic antimicrobial HDPs offer a significantly wider therapeutic window than their lytic counterparts^12,13^. As exemplified by the proline-rich HDPs (PrHDPs) family, this translocation is frequently facilitated by specific inner-membrane (IM) transporters (e.g., SbmA and the MdtM-YjiL cluster^12,14–16^). Crucially, because mammalian cells lack these transporters, they provide an intrinsic safety layer enabling lower effective dosing and minimal partitioning into host tissue.

Nevertheless, reliance on specific IM transporters narrows the activity spectrum of PrHDPs^5,6,11^. Although certain PrHDPs like PR-39^17^ exhibit activity against Gram-positives, most family members target only those Gram-negatives that do express compatible IM transporters. This reliance also creates a vulnerability to resistance by transporter loss. However, combining orthogonal uptake pathways of non-lytic HDPs can potentially overcome this limitation, broaden the activity spectrum and enhance cellular uptake^18,19^. In fact, because peptides within the wider non-lytic HDP group also bind to diverse intracellular machinery, fusing distinct non-lytic HDPs into a single macromolecule can potentially yield multimodal chimeras that inherit these complementary properties.

While multimodal molecular designs are heavily exploited in oncology^20^ and targeted drug delivery^21^, they have remained mostly underexplored in the field of antimicrobial HDPs. Previous research on chimeric HDPs has identified a few candidates with dual MOAs and potentiated activity^22–25^, but these efforts have been restricted to small, chemically synthesized libraries (fewer than 20 variants) primarily due to prohibitive bottlenecks such as synthesis costs and technical screening constraints. Furthermore, rational design remains hindered by a poor understanding of HDP sequence-activity relationships (SAR) as the residues involved in cellular uptake or target binding are frequently unknown, thereby diminishing the predictability of the functionality of *in silico* designed HDP chimeras. Consequently, a comprehensive implementation of HDP chimerization remains missing, specifically one that deploys diverse design principles and leverages structural variation to compensate for lack of functional insight.

We addressed these bottlenecks by implementing a high-throughput framework to explore the large activity landscape of chimeric non-lytic HDPs. We constructed a gene library of 99,235 “chimerophores” by encoding the systematic translational fusion of 56 non-lytic parent HDPs via a diverse repertoire of amino acid linkers and in a variety of configurations into DNA. To assay chimerophore function, we utilized the Me^x^ workflow^26^, relying on next-generation sequencing (NGS) to assess growth inhibition upon ribosomal synthesis of each variant in *Escherichia coli*. This approach bypasses the chemical peptide synthesis bottleneck and enables the primary activity screening of nearly 100,000 unique chimerophores.

The Me^x^ assay allowed for the selection of a promising subset of chimerophores for in-depth analysis. Remarkably, the selected chimerophores showed a broader spectrum of activity and higher serum tolerance than their parent HDPs. We confirmed their multimodal MOAs and synergistic potencies against ESKAPE pathogens, validating the potential of this engineering approach. Finally, the selected chimerophores retained the favorable *in vitro* safety profiles of their parents. Notably, further evaluation of a lead chimerophore revealed that it significantly suppressed the emergence of resistance in *Pseudomonas aeruginosa*. Given this pathogen’s high intrinsic resistance and propensity to rapidly mutate under single-target selective pressure^27^, this finding demonstrates that multimodal chimerophores can successfully mitigate mutational escape. Overall, our findings establish a scalable platform for the systematic study of chimeric peptides, offering a promising path forward for the clinical translation of ribosomally synthesized non-lytic HDPs as durable antimicrobial therapies.

## Results

### Selection of the parent HDP set

Aiming to engineer multimodal antimicrobials, we designed a DNA-encoded library of 99,235 chimerophores from intracellularly targeting parent HDPs in diverse configurations. We first selected 12 natural HDPs from diverse species (Fig. 1a; Extended Data Table 1; Supplementary Fig. 1) exhibiting potent antimicrobial activity against pathogens of the ESKAPE group and low toxicity to mammalian cells^12,13^. These HDPs included PrHDPs, (e.g., apidaecin 1B (api1B), oncocin (onc), and drosocin (dro)) typically adopting extended polyproline II helical conformations and targeting 70S ribosomes. We also included intrinsically “disordered” non-PrHDPs (e.g., Trp-rich indolicidin (indo) and the Arg-rich buforin 2 (buf2)), which remain disordered in solution but adopt secondary structures upon contact with membranes and preferentially target nucleic acids. Additionally, we included 44 engineered analogs derived from these 12 natural HDPs, either sourced from the literature or designed in this work with targeted amino acid substitutions. This resulted in a set of 56 non-lytic parent HDPs (Supplementary Table 1).

**Figure 1.**
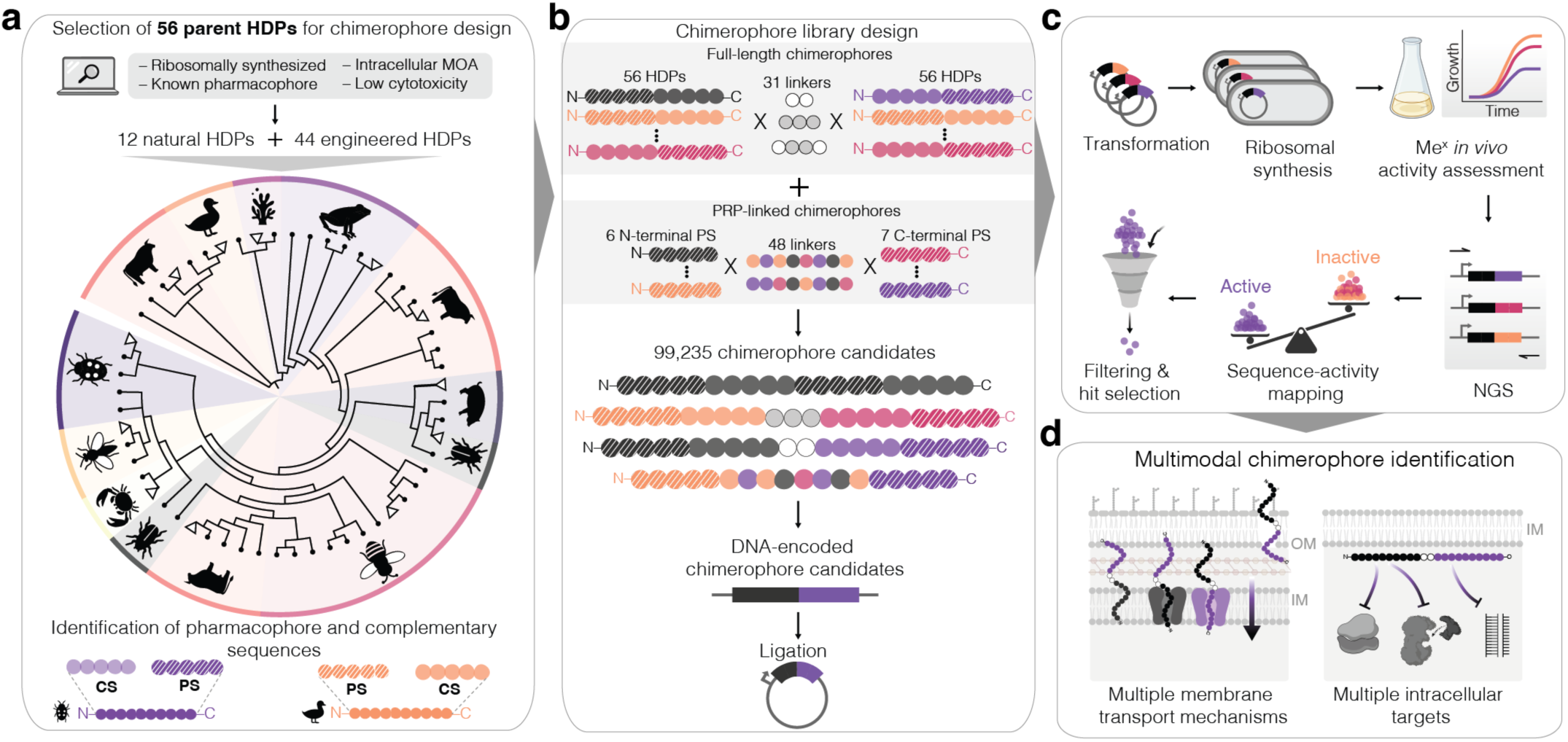
Chimerophore design and discovery. **(a)** Selection of parent host defense peptides (HDPs) for design of a library encoding putatively multimodal chimerophores. The species diversity of the set of 56 parent HDPs is represented in a dendrogram, which organizes 12 natural non-lytic HDPs (triangle) along with 44 engineered analogs (black circles) sourced from literature or designed in this work. These parent HDPs originate from 10 species (diverse background shading colors), listed clockwise: *Ciona intestinalis, Bufo bufo gargarizans, Bos taurus, Sus scrofa, Oncopeltus fasciatus, Apis mellifera, Hyas araneus, Drosophila melanogaster, Pyrrhocoris apterus,* and *Anas platyrhynchos*. The pharmacophore sequences (PS) containing essential residues for intracellular target binding (lined circles) and complementary sequences (CS) containing residues that mediate transport through bacterial membranes (solid circles), were identified in the literature prior to generation of the library. **(b)** Chimerophore candidate-encoding DNA library construction resulting in 99,235 variants. Full-length chimerophores were designed by combining the 56 parents in pairs, connected directly or via 2-4mer linkers consisting of small flexible/neutral (G, S, T; white circles) or basic amino acids (R, K, H; gray circles). For PRP-linked chimerophores, six N-terminal PS were connected to seven C-terminal PS via 11mer linkers (mixed colors) derived from conserved motifs in the CS of parent HDPs. The DNA-encoded chimerophore library was inserted in plasmid vectors for arabinose-inducible expression in *Escherichia coli* strains. **(c)** High-throughput screening of the chimerophore library using the parallelized growth assay Me^x^. *E. coli* cells expressing antibiotic chimerophores are growth-inhibited, while cells expressing inactive peptides continue to grow. Chimerophore-encoding DNA abundance is quantified over time by NGS enabling the discrimination between putative antibiotic chimerophores and inactive ones. Active variants were filtered based on favorable physicochemical properties (hydrophilicity, peptide size, positive net charge and instability index) to select a few variants for chemical synthesis and further characterization. **(d)** This chimerophore design potentially yields multimodal chimerophores that employ more than one cell entry mechanism to traverse the outer (OM) and inner membrane (IM) of Gram-negative bacteria and once inside the cell act on multiple essential targets (e.g., 70S ribosome and nucleic acids).

### Chimerophore design

We designed two chimeric architectures: full-length and PRP-linked chimerophores. Most of the library comprised full-length chimerophores (97,216 variants), in which two parent HDPs were systematically fused in all possible pairwise combinations (i.e., A-B, B-A, A-A, and B-B) (Fig. 1b). Coupling was achieved either directly (56^2^ = 3,136 variants) or via 11 “flexible” linkers consisting of small, neutral residues (e.g., GS, GTGT; 34,496 variants) or 19 “basic” linkers being cationic at neutral pH as they were enriched with Arg, Lys, or His (e.g., RR, GRRG; 59,584 variants). Notably, basic linkers contain cleavage sites for the *E. coli* endopeptidase OpdB, preferentially cleaving peptides of up to 40 amino acids^28^ upstream of basic residues^26^. We hypothesized that OpdB cleavage could intracellularly release the parent modules from chimerophores that may obstruct macromolecular target binding while fused.

For the PRP-linked chimerophore library (2,016 variants), we conceptualized each parent HDP as a modular construct containing two partially overlapping functional segments based on established SAR^29–33^. The first is the pharmacophore sequence containing the essential target-binding amino acids, and the second is the remaining sequence with amino acids often involved in cellular uptake^14,15,34,35^. We fused the pharmacophore sequences of 13 parent HDPs retaining their natural positioning at the N-termini (for 6 HDPs, e.g., bac5 and bac7) or C-termini (for 7 HDPs, e.g., api1B and dro) (Extended Data Fig. 1a). These were connected via one of 48 distinct 11-mer linkers derived from a consensus sequence generated by aligning the remaining sequences of the 13 parent HDPs (Extended Data Fig. 1b; see Methods). These linkers incorporate two central PRP motifs aiming to promote structural stability via polyproline II helices, while maintaining affinity for IM transporters^14,15^. This yielded a library of 99,235 chimerophores, including three poly-Gly peptide variants used as negative controls (Supplementary Table 2).

### Chimerophore activity landscape

We transformed *E. coli* cells with the chimerophore-coding DNA sequences and used Me^x^ for initial activity testing (Fig. 1c) and identification of candidates for subsequent characterization (Fig. 1d). Me^x^ tracks the relative frequency of each chimerophore-encoding gene within the population over time, whereby sequence depletion upon chimerophore-expression indicates antimicrobial activity (Extended Data Fig. 2a).

Importantly, antimicrobial activity is measured in a cellular environment, in which defense mechanisms such as efflux pumps and host proteases are active. Therefore, intrinsically inactive, rapidly secreted, or degraded variants are deselected.

At the time of induction t_0_ (Extended Data Fig. 2b), we detected 99,047 (99.8% of all designed) chimerophores, with 75,196 (75.9%) meeting the detection frequency thresholds for robust differential abundance analysis. We binned active and inactive chimerophores into five activity categories based on their log_2_-fold change (LFC; Fig. 2b), ranging from highest growth-inhibiting activity (category 1; LFC < -6) to no significant activity (category 5; -1 < LFC < 0).

**Figure 2.**
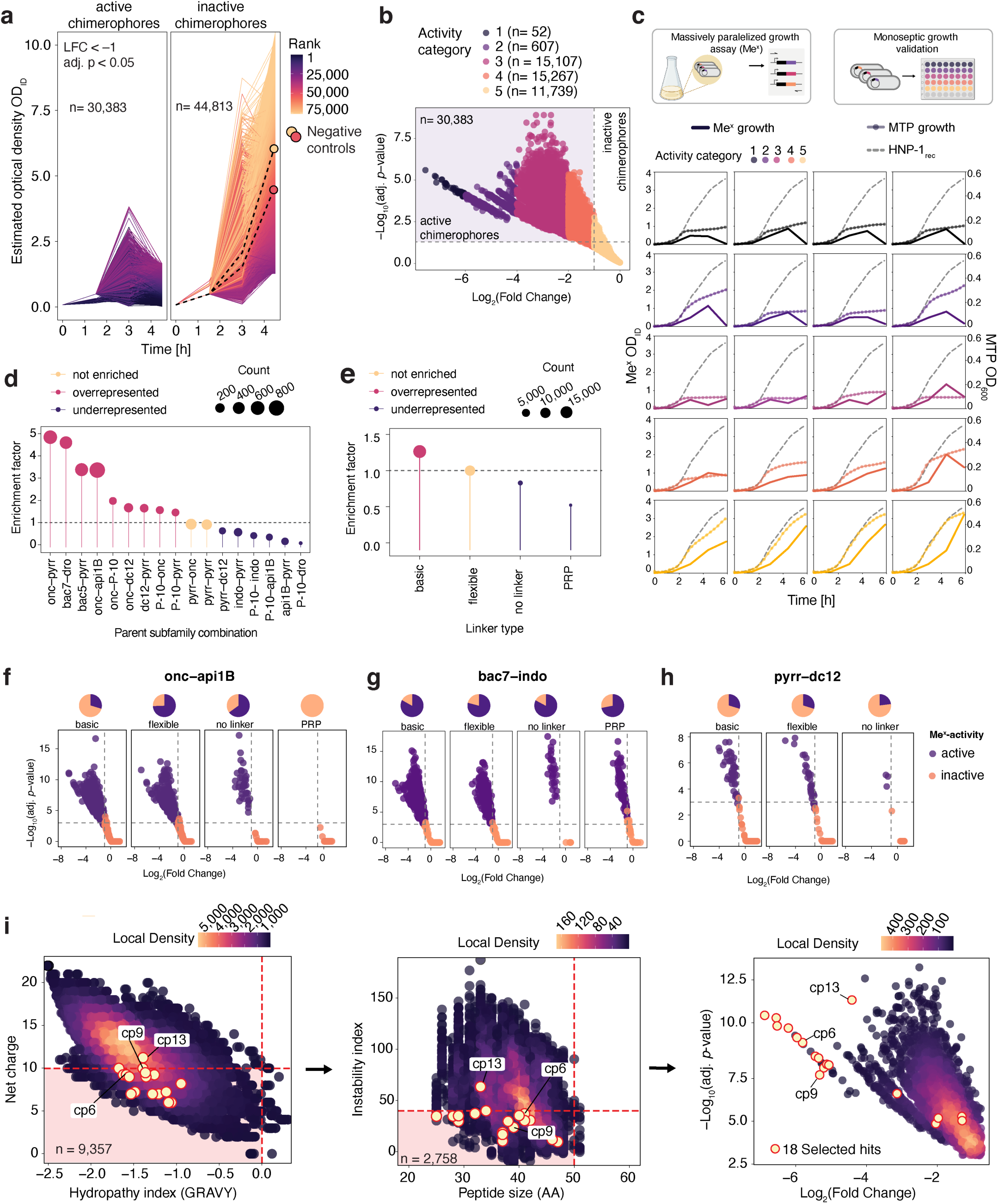
Me^x^ screening, enrichment analysis and selection of top active chimerophores. **(a)** Growth curves (OD_ID_) estimated from the differential abundance of chimerophore-encoding DNA in a Me^x^ culture of *E. coli* strains carrying a plasmid library of putatively active chimerophores. Activity thresholds for distinguishing between active and inactive chimerophores were set at Log_2_(fold change) (LFC) of chimerophore-encoding DNA counts below –1 and adjusted p-value < 0.05 (One-sided Wald’s test, Benjamini-Hochberg correction). Variant with the lowest OD_ID_ at t_3_=4.5 h is the highest-ranking. Negative controls are *E. coli* strains expressing either a 38mer poly-Gly or a 38mer poly-Gly-Ser sequence (dashed black lines). The data set consists of growth curves for strains expressing n=75,196 chimerophore sequences for which we obtained sufficient reads at the time of chimerophore expression induction t_0_=0 h (NGS read count ≥15). **(b)** Chimerophores sorted into activity categories based on LFC values in a one-tailed volcano plot (activity thresholds indicated as black dashed lines), the quadrant with active chimerophores indicated in purple shading. The categories are defined by LFC thresholds: category 1 (strongest inhibition): LFC <−6; category 2: −6≤ LFC <−4; category 3: −4≤ LFC <−2; category 4: −2≤ LFC <−1; and category 5: –1≤ LFC <0. Inactive candidates (LFC ≥0) are not shown. **(c)** Validation of growth behavior of four randomly selected strains per activity category from the Me^x^ assay in monoseptic culture in microtiter plates (MTPs). Growth was followed by turbidity measurements over time (dot and line) and overlayed with the OD_ID_ curves calculated in Me^x^ (solid line) based on chimerophore-encoding DNA NGS read counts. The recombinantly expressed HNP-1_rec_ (dashed line) was used as an inactive HDP control. Data points are calculated from n=3 biological replicates. **(d)** Enrichment and depletion analysis for a representative subset of parent subfamily combinations in the Me^x^-active fraction, calculated via a two-sided Fisher’s exact test with Benjamini-Hochberg correction for multiple testing, and adjusted p-value cut-off <0.05. **(e)** Differential representation of linker types in the Me^x^-active chimerophore fraction; PRP: linkers containing Pro-Arg-Pro motifs. **(f-h)** Representative activity profiles for three parent subfamily combinations grouped by linker type. Pie charts illustrate the proportions of active (purple) and inactive (orange) chimerophores derived from two significantly enriched parent subfamily combinations onc–api1B (f) and bac7–indo (g), and from the significantly underrepresented combination pyrr–dc12 (h); activity thresholds indicated by gray dashed lines. **(i)** Filtering process for the selection of 18 chimerophores (red circles with yellow fill) for chemical synthesis and further characterization. Dashed lines represent the distinct physicochemical property filters applied (quadrant area shaded in pink): positive net charge < 11, hydropathy index (GRAVY) < 0, instability index < 41 and peptide size < 51. Selected chimerophores cp6, cp9 and cp13 are labelled as examples throughout the filtering process.

We next tested whether similar results would have been obtained if the screening strains had been cultivated axenically. For this, we evaluated the growth of four randomly selected strains from each Me^x^ activity category in microtiter plates. An overlay with the Me^x^ data indicated strong growth inhibition in categories 1 to 4, but only a modest inhibitory effect in category 5 (Fig. 2c). This confirmed that the Me^x^ results remained vastly unbiased by phenomena such as nutrient competition, peptide leaching, or intracellular signaling^26^, thereby validating the statistical approach based on LFC as a proxy for activity. Ultimately, a surprisingly large proportion of 30,383 (40.4%) fell into activity categories 1 to 4 and were classified as “Me^x^-active” (Fig. 2a).

Next, we analyzed the entire Me^x^-data set using Fisher’s exact tests to identify major trends and design rules. The relative frequency at which chimerophores were Me^x^-active was highest (53%) for fusions of two PrHDPs (Extended Data Fig. 3a, b). Substitution with disordered non-PrHDPs at the N- or C-terminus reduced the active fraction to 28% and 42%, respectively, and to 9% for dual non-PrHDP combinations. These class-specific differences are plausibly due to chimerophore proteolytic degradation by exopeptidases occurring concurrently with translation.

Assuming that membrane transport and target binding are common among structurally similar parent HDPs, we analyzed the dataset by subfamily, each centered on one of the 12 wild-type HDPs (e.g., api1B_WT_) and their engineered analogs (e.g., api1B_G_ and api1B_RVR_). The parent subfamilies, bac5, bac7, and onc yielded the highest ratios of proven active-to-inactive chimerophores in Me^x^ (Extended Data Fig. 2c). For example, over 79% (2,400 out of n=2,964 used in differential abundance analysis) of the evaluated variants containing bac7_PSL_, onc_KR_, or bac5_G_ were active in Me^x^. Conversely, buf2, pyrr, and dro showed the lowest activity ratios. For instance, buf2_G,_ pyrr_G_, and dro_4_ yielded only 15% (400) Me^x^-active variants, even though at least 80% of their respective total designed sequences (n=3,472) were included in the differential abundance analysis.

Generally, PrHDP-derived chimerophores, such as onc–pyrr, bac7–dro, and onc–api1B, were considerably overrepresented in the Me^x^-active set (Fig. 2d; Extended Data Fig. 3c). This preference was highly directional; for instance, onc–pyrr variants yielded 75% actives while the pyrr–onc orientation only 38% (Fisher’s exact test, adjusted p > 0.05). Similar disparities were observed for bac7/dro (74% vs. 11%) and onc/api1B combinations (68% vs. 28%) (Extended Data Fig. 3d). These results likely reflect the specific target binding requirements of the parental HDPs as onc, bac7 and pyrr (Type I translation inhibitors) enter the ribosomal exit tunnel via their N-terminus^36–38^, but dro and api1B (Type II translation inhibitors) via their C-terminus^39,40^. Extension of these termini upon coupling to another HDP therefore likely abolishes activity, explaining why chimerophores comprised of the same parent HDP modules but in reversed configurations fail to inhibit growth in Me^x^.

Similar observations were made in other subfamilies remaining Me^x^-inactive in one but not the reverse orientation (Extended Data Fig. 2d). For instance, PrHDPs ara1, api1B, and dro fused to the N-terminus of the Lys-rich disordered peptide P-10 were generally Me^x^-inactive. These results align well with established PrHDP-ribosome interactions. Dro and api1B are Type II ribosomal inhibitors, and while the ara1 binding site is unknown, all three likely require a free C-terminus to entering the ribosomal exit tunnel^39^. Similarly, chimerophores containing buf2 at the N-terminus were inactive in Me^x^ when fused to ara1, dro, or any of the disordered HDPs P-10, dc12, indo, or to itself. This suggests that C-terminal fusions of those HDPs to buf2 abolish its activity^41^ and that the respective partners could not rescue activity.

Fisher’s exact test further revealed that the linker type influences chimerophore function. Relative to the whole library, chimerophores with basic linkers were significantly overrepresented in the Me^x^-active set (42% active) but not those with flexible linkers (40% active) (Fig. 2e). Conversely, PRP- or no-linker chimerophore variants were underrepresented (26% and 35% active, respectively) (Fig. 2e), suggesting that basic or flexible linkers generally promoted activity in Me^x^. Within this trend, activity varied considerably depending on the specific pairing and positioning of two parent subfamilies. For example, the subfamily combinations onc–api1B (68% active) and pyrr–dc12 (29% active) both mirrored the general linker preferences (Fig. 2f, h). However, onc–api1B could be coupled directly without a linker (65% active) but failed with PRP linkers (0% active). Notably, the bac7–indo subfamily combination deviated significantly, maintaining high success rates (>70% active) across all linker types, including the generally unfavorable no-linker (83% active) and PRP-linker variants (72% active) (Fig. 2g).

Additionally, basic linkers were essential for chimerophore activity in the subfamily combinations api1B–dc12, indo–P-10, ara1–dc12, ara1–dro, dro–dro, api1B–ara1, ara1–ara1, and P-10–P-10 (Extended Data Fig. 4). This preference was unique to basic linkers, as no other linker type rescued activity in otherwise inactive parent pairings. These observations highlight how minor structural changes modulate chimerophore efficacy against intracellular targets.

Unexpectedly, we observed growth-inhibiting activity in parent fusions in blocked configurations. Specifically, in api1B–bac5 chimerophores (including all subfamily members), where the C-terminus of the Type II ribosome-inhibiting parent (api1B) is covalently linked to the N-terminus of the Type I inhibitor (bac5). This design was expected to preclude chimerophore access to the ribosomal exit tunnel but remarkably, 45 out of the 50 combinations still elicited a strong Me^x^-response. Activity was linker-independent in 39 of these cases, while five required a basic linker (Supplementary Fig. 2). As the remaining six structurally similar subfamily combinations remained inactive, these results point toward target specificity and likely the activation of ribosome-specific pathways, perhaps involving endoproteolytic cleavage enabling release of the structurally buried termini.

Taken together, the Me^x^ results yielded the following design guidelines: (i) PrHDPs are promiscuous fusion partners at either terminus; (ii) non-PrHDPs should be placed at the C-terminus; and (iii) basic linkers are favorable particularly in full-length chimerophore designs. Beyond these trends, specific parent subfamily compatibility remains highly case-dependent, confirming the utility of a comprehensive screen for navigating the non-intuitive activity landscape of chimeric HDPs.

### Selection of chimerophores for further characterization

We next sought to characterize a representative subset of chimerophores if administered extracellularly to bacteria. For this, we applied physicochemical thresholds facilitating solid phase peptide synthesis and purification^42^ as selection criteria (Fig. 2i; net charge < +11; negative GRAVY score^43^; <51 amino acids in length)^42^. In addition, an instability index <41 was set for high predicted stability *in vitro* and potentially extended half-lives *in vivo*^44^. This produced a subset of 2,758 Me^x^-active chimerophores (Fig. 2i) from which 18 were synthesized. Of these, half were dual PrHDPs and half were PrHDP–non-PrHDP combinations. We argue that these 18 Me^x^-active chimerophores comprise a representative and diverse set of HDP subfamilies, their pertaining MOAs, and species of origin (Extended Data Table 2; Supplementary Fig. 3).

### Chimerophores can have a broader spectrum of activity than parent HDPs

Remarkably, all 18 selected chimerophores demonstrated potent activity with low micromolar minimal inhibitory concentrations (MIC) and eradicated at least two Gram-negative ESKAPE pathogens in Mueller Hinton Broth (MHB) (Fig. 3a). Notably, they integrated and expanded the activity spectrum of their parents. For instance, cp14, a PRP-linked chimerophore derived from disordered non-PrHDPs (P-10_WT_ and indo_WT_), exhibited a 14-fold and 4-fold lower MIC than its parents against *P. aeruginosa* and *Staphylococcus aureus*, respectively, and gained activity against the Gram-negative *Klebsiella pneumoniae*, a species not targeted by either parent (Fig. 3a). Furthermore, while PrHDP parent activity is generally restricted to Gram-negative bacteria, 11 chimerophores containing at least one PrHDP demonstrated potency against Gram-positive *S. aureus*, even when neither parent showed activity on its own (Fig. 3a). For instance, while the Trp-rich parent dc12_G_ was inactive for all ESKAPE strains, the dc12_G_-containing cp2 not only preserved the anti-Gram-negative spectrum of its PrHDP parent onc_KR_ but also expanded if to include *S. aureus*. Similarly, cp6 expanded the spectrum of its anti-Gram-negative PrHDP parents (onc_KR_ and api1B_G_) to target *S. aureus*. These findings demonstrate that this chimeric design offers a robust route to convert narrow spectrum HDPs into broad-spectrum chimerophores. We next selected the 12 most potent, broad-spectrum chimerophores for further characterization.

**Figure 3.**
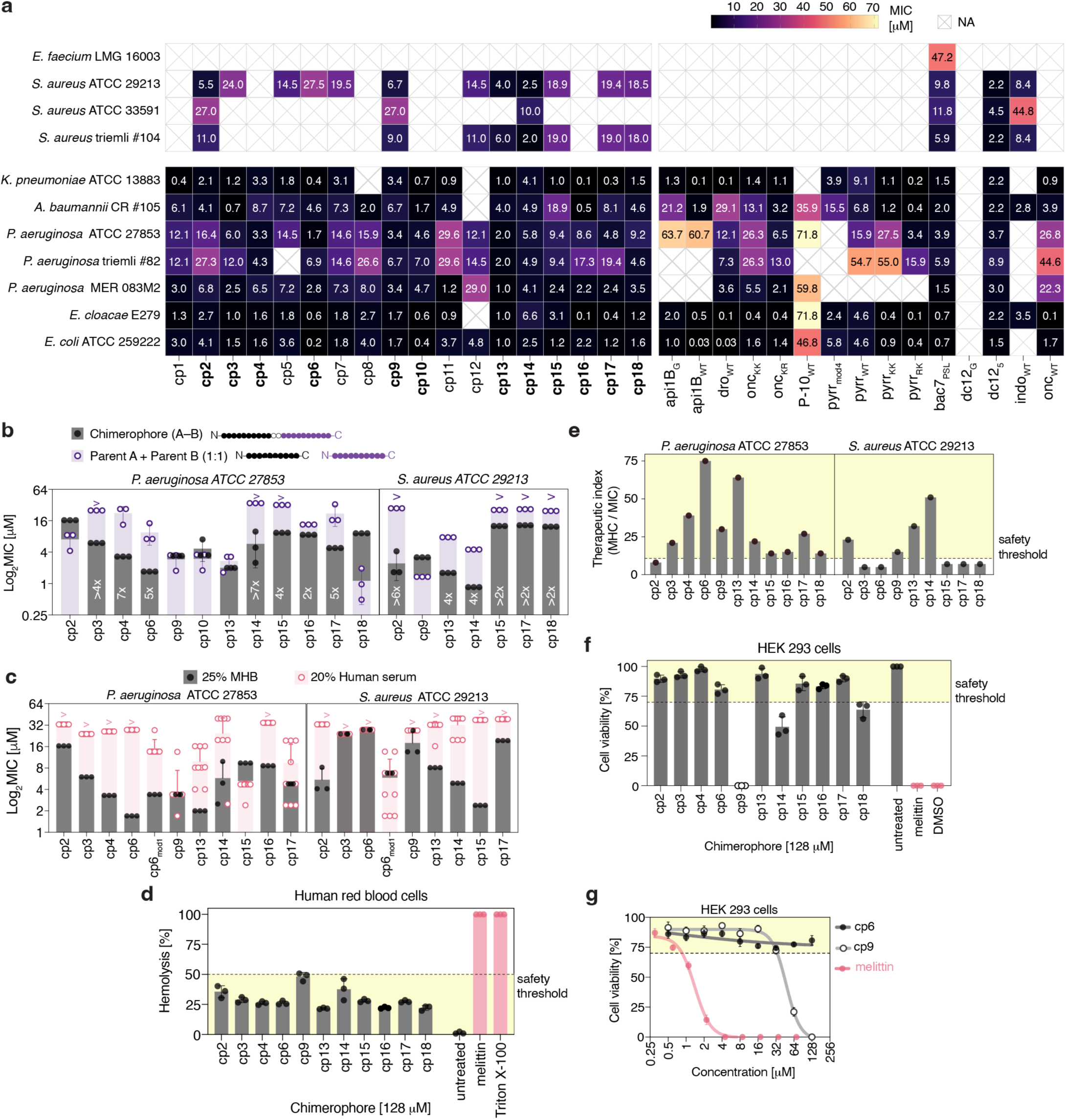
Antibiotic activity and toxicity profiling of selected chimerophores. **(a)** MIC values of chemically synthesized chimerophores cp1 to cp18 (left) and the 15 parent HDPs used for their construction (right) tested against ESKAPE bacterial pathogens including Gram-positives (*Enterococcus faecium*, *Staphylococcus aureus*) and Gram-negatives (*Klebsiella pneumoniae*, *Acinetobacter baumannii*, *Pseudomonas aeruginosa*, *Enterobacter cloacae*, and *Escherichia coli*) in 25% MHB. The mean of n=3 biological replicates is shown. NA (not applicable, white crossed boxes): MIC values greater than the highest concentration tested indicating no inhibition up to that concentration. A subset of 12 chimerophores selected for further assessment are indicated in bold. **(b)** MIC assays in 25% MHB comparing antimicrobial activity of the 12 chimerophores selected in (a) (black) with MIC below 10 µM against *P. aeruginosa* or *S. aureus*, and their parent HDPs in 1:1 molar mixture (purple) against *P. aeruginosa* ATCC 27853 (left) and *S. aureus* ATCC 29213 (right). Bars represent the mean of n=3 biological replicates, error bars represent the standard error of the mean. “>” indicates undetermined MIC at the highest concentration tested. **(c)** Antibacterial activity of 10 chimerophores selected from (b) assessed in MIC assays against *P. aeruginosa* ATCC 27853 in 20% human serum (pink open circles) or 25% MHB (black circles, data from (b)). Bars represent the mean and error bars the standard deviation of n=3 biological replicates for 25% MHB and n=6 for 20% human serum MIC measurements. “>” indicates an undetermined MIC at the highest concentration tested. **(d)** Hemolytic effect of chimerophores at 128 µM (black circles) compared to the highly hemolytic HDP melittin (pink circles) and Triton X-100 on human red blood cells recorded in MTPs. Acceptable hemolysis level cut-off at highest concentration tested was set to <50% (area shaded in yellow). Columns and error bars represent the mean and standard deviation of n=3 biological replicates. **(e)** Therapeutic index (TI) of chimerophores (black circles) calculated as the ratio between the minimum hemolytic concentration (MHC) and the MIC for *P. aeruginosa* ATCC 27853 (left) and *S. aureus* ATCC 29213 (right). A favorable TI cut-off was set to >10 (area shaded in yellow). Columns and error bars represent the mean and standard deviation of n=3 biological replicates. **(f)** *In vitro* assessment of cytotoxicity on HEK 293 cells. Chimerophore treatment at 128 µM (black) is compared to the positive control melittin (pink), which is unsuitable for human treatment due to its high cytotoxicity. Acceptable cytotoxicity cut-off was set to >70% (area shaded in yellow). Columns and error bars represent the mean and standard deviation of n=3 biological replicates. **(g)** Effect of chimerophore concentration on cell viability: comparison of dose-cell viability response curve of the membranolytic HDP melittin (pink circles), cp6 (black circles), and cp9 (open black circles) in HEK 293 cells. Nonlinear regression fit with confidence level 95%, dots represent the mean of n=3 biological replicates, error bars represent the standard deviation of the mean. Dashed line indicates the cytocompatibility cut-off at >70% cell viability (area shaded in yellow).

We assessed chimerophore potency by comparing their MICs against *P. aeruginosa* and *S. aureus* to that of the parent equimolar mixtures (Fig. 3b). In 11 out of 12 cases, chimerophore potency was equal to or greater than (synergistic effect) their parent mixture. For example, cp14 and cp6 demonstrated synergistic activity against *P. aeruginosa*, with MICs 8-fold and 4-fold lower than their parental equimolar mixtures. Similarly, cp2, cp15, cp17 and cp18 displayed synergistic activity against *S. aureus*, with MICs at least 2- to 6-fold lower than their inactive parent mixtures (MICs >128 µg mL^-1^). Collectively, these results demonstrate that the covalent linkage of two parent HDPs enables intramolecular synergy, yielding chimerophores significantly more potent than their unlinked parents.

### Chimerophores are active in serum and show low cytotoxicity

Clinical translation of HDPs is frequently hampered by a loss of activity in human serum, resulting from proteolysis or from divalent cations (e.g., Mg^2+^ and Ca^2+^) shielding negative charges on the bacterial surface and blocking peptide uptake^5^. We tested chimerophore activity against *P. aeruginosa* and *S. aureus* in 20% human serum to mimic physiologically relevant conditions while permitting optical density-based MIC determination^45^.

Chimerophores cp9, cp13, cp14, cp15 and cp17 lost activity against *S. aureus* but remained active against *P. aeruginosa* in 20% serum (Fig. 3c). Interestingly, the activity of parents dc12_5_ and bac7_PSL_ against *P. aeruginosa* also remained unaffected (Supplementary Table 3), a feature that seems to be inherited by their derived chimerophores cp9 and cp13, respectively. Conversely, the PRP-linked chimerophores cp14, cp15, and cp17 outperformed their parents, P-10_WT_, indo_WT_, dro_WT_, and api1B_WT_, which uniformly lost activity in serum (Fig. 3c; Supplementary Table 3). These results indicate enhanced serum tolerance of chimerophores relative to their parents perhaps due to reduced susceptibility to proteolytic degradation or salt-induced inactivation, facilitated by modulated physicochemical properties or enhanced structural stability. Supporting this, AlphaFold modeling of cp9 predicted an intramolecularly stabilized fold that may shield potential cleavage sites from proteases (Supplementary Fig. 3).

Nevertheless, many full-length chimerophores with broad-spectrum activity in MHB, such as cp2 and cp6, lost activity in serum against both *P. aeruginosa* and *S. aureus* (MIC > 128 µg mL^-1^). To restore cp6 activity, we introduced the substitutions Arg15/D-Arg and Arg19/D-Arg in the onc module and Gly1/Orn in the api1B module (cp6_mod1_), as these modifications are known to increase serum tolerance in the individual parents via enhanced proteolytic stability^39,46,47^. In serum, cp6_mod1_ exhibited a partial recovery of activity against *P. aeruginosa*, with 4-fold higher MIC compared to MHB. However, its activity against *S. aureus* could be rescued with an MIC 4-fold lower than that of cp6 (Fig. 3c). Targeted chemical modifications can therefore be exploited to enhance activity in serum, positioning chimerophores as excellent scaffolds for optimization.

To determine if chimerophores maintained the low hemolysis and cytotoxicity of their parents, we exposed human red blood cells and HEK 293 cells to increasing peptide concentrations. All chimerophores showed less than 50% hemolysis at up to 128 µM (Fig. 3d) (Extended Data Fig. 5), therefore yielding undetermined minimal hemolytic concentrations (MHC_50_>128 µM). We next assessed the therapeutic index (TI) as the ratio of the hemolytic dose (MHC_50_) to the respective MIC values, considering TI values above 10 as safe^48,49^. Importantly, the TI of most chimerophores exceeded the threshold and was consistently equal to or greater than that of their parents. For instance, cp4, cp6, cp9 and cp13 against *P. aeruginosa*, and cp2, cp13 and cp14 against *S. aureus*, exhibited TI values greater than 20 (Fig. 3e). Moreover, cp6 achieved a TI of 75, by far exceed those of its parents (onc_KR_, TI=19.7; api1B_G_, TI=2). These results confirm that this chimeric design strategy can preserve or enhance the favorable TIs of non-lytic parent HDPs.

To evaluate the *in vitro* cytotoxicity profile of chimerophores, we defined cytocompatibility as greater than 70% cell viability at 20-fold the respective MIC against *P. aeruginosa*^50^. Assessing cell viability in HEK 293 cells, we found that many chimerophores met this requirement (Fig. 3f, Extended Data Fig. 6). For example, cp6 showed a flat dose-response curve within the safety threshold, with an undetermined IC_50_ even at the highest concentration tested (128 µM) (Fig. 3g), whereas the lytic control HDP melittin drastically reduced cell viability at 4 µM (less than two-fold its MIC). While cell viability for cp9 dropped below the 70% safety threshold at 128 µM, this concentration is more than 37-fold its MIC against *P. aeruginosa* and therefore still indicates cytocompatibility. Collectively, these findings demonstrate that the 18 chimerophores successfully retained the low cytotoxicity of their non-lytic parents, validating our design strategy as a reliable route to engineer safe antimicrobials with high bacterial cell selectivity.

### Chimerophores retain the non-lytic features of their parent HDPs

To confirm whether chimerophores maintain non-lytic MOAs at the MIC, we assessed membrane damage in *E. coli* by flow cytometry^51^ using propidium iodide uptake (PI) and GFP retention as markers (Fig. 4a). We gated cells after peptide treatment into four subpopulations: intact membrane (GFP+/PI-), small-pore formation (GFP+/PI+), macro-pore formation (GFP-/PI+), and membrane rupture accompanied by DNA loss (GFP-/PI-) (Extended Data Fig. 7a-d). Unlike the membranolytic control LL-37, most chimerophores (except cp14) were non-membranolytic at the MIC, consistently showing greater than 70% of cells with intact membranes (GFP+/PI-) (Fig. 4b, Supplementary Fig. 4) and aligning with the profiles of their parental HDPs. These results confirm that indeed 17 out of the 18 chimerophores traverse the bacterial envelope of *E. coli* and exert activity without membrane disruption.

**Figure 4.**
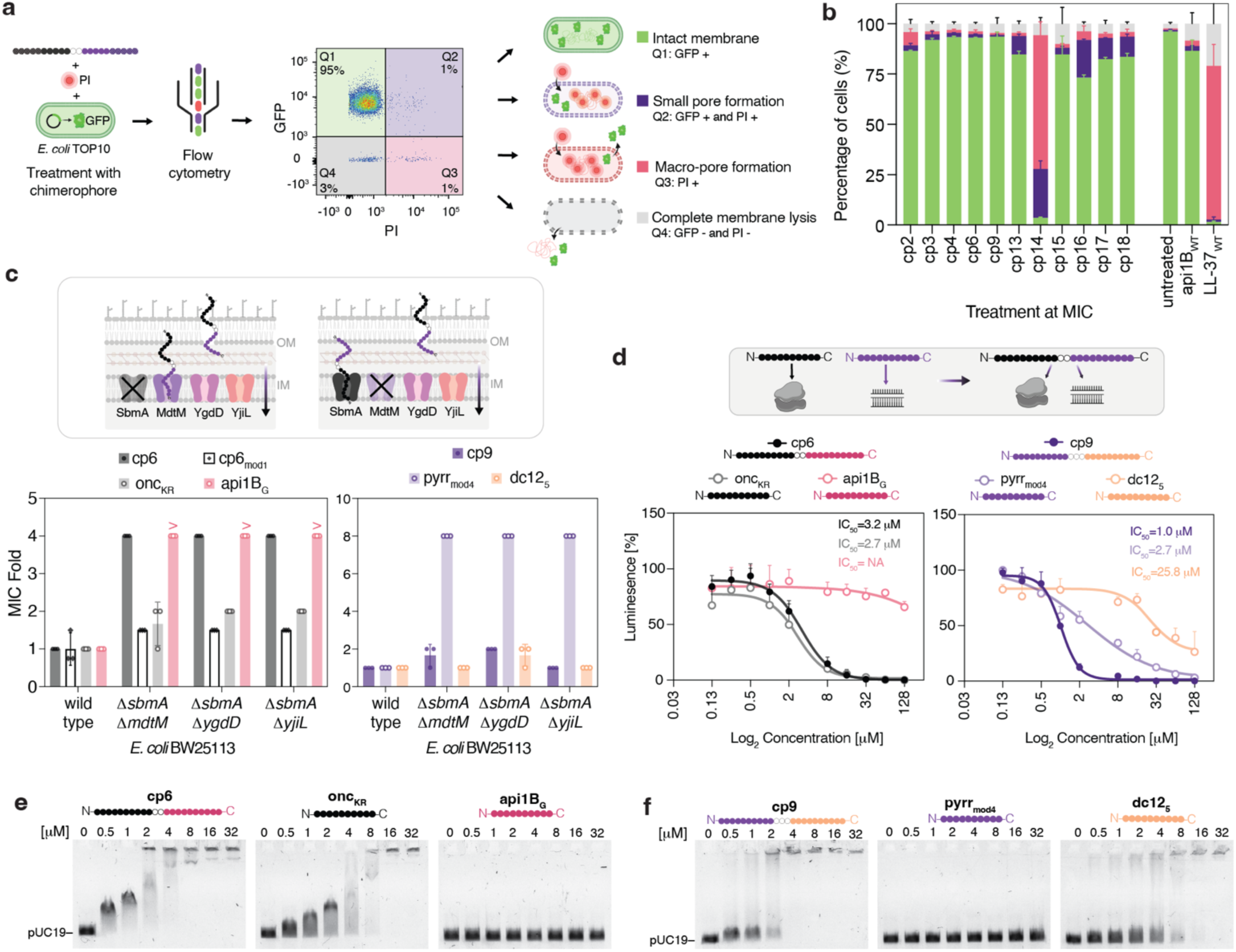
Evaluation of the MOA of selected chimerophores in Gram-negative bacteria. **(a)** Workflow for assessment of the membrane status of *E. coli* TOP10 constitutively expressing a green fluorescent protein (GFP) by flow cytometry. GFP+ indicates an intact bacterial membrane (green), GFP+ PI+ indicates the formation of small pores allowing PI to enter the cells and stain the genomic DNA (purple), PI+ indicates macro-pore formation leading to massive loss of GFP (pink), and GFP- PI-indicates complete membrane rupture leading to the washout of DNA and GFP, therefore loss of both PI and GFP fluorescence (gray). **(b)** Membrane damage by chimerophores (left) and controls (right) including the non-membranolytic parent PrHDP api1B_WT_ and the membranolytic HDP LL-37 (used as reference but not for library design) in *E. coli* TOP10 cells constitutively expressing GFP. Bars represent the mean of n=2 biological replicates, and error bars represent the standard deviation. **(c)** Activity assays against double knockout strains *E. coli* BW25113 *ΔsbmA ΔmdtM*, *E. coli* BW25113 *ΔsbmA ΔygdD,* and *E. coli* BW25113 *ΔsbmA ΔyjiL*. Each double knockout removes two IM transporters utilized by PrHDP parents for cellular uptake. Cells were treated with chimerophores cp6 (solid filled black circles), cp9 (solid filled purple circles) and their parent HDPs (open circles) onc_KR_ (gray, PrHDP), api1B_G_ (pink, PrHDP), pyrr_mod4_ (transparent purple, PrHDP), and dc12_5_ (orange, Trp-rich HDP). The MIC-fold is calculated relative to the MIC values in wild type *E. coli* BW25113. Columns represent the mean and error bars the standard deviation (n=3 biological replicates). “>” indicates undetermined MIC at the highest concentration tested. **(d)** Inhibitory activity of increasing concentrations of chimerophores and their parent HDPs on *in vitro* transcription/translation using the firefly luciferase (Fluc) as reporter. To the left, cp6 (black) and its parent HDPs onc_KR_ (gray) and api1B_WT_ (pink). To the right, cp9 (purple) and its parent HDPs pyrr_mod4_ (transparent purple) and dc12_5_ (orange). Nonlinear regression fit with confidence level 95%, dots represent the mean of n=3 biological replicates, error bars represent the standard deviation of the mean. **(e)** Electrophoretic mobility shift assay monitoring the nucleic-acid binding ability of increasing concentrations of chimerophore cp6 (left) and its parent HDPs (onc_KR_ middle, api1B_G_ right). Linearized pUC19 plasmid DNA was used. **(f)** Electrophoretic mobility shift assay monitoring the nucleic-acid binding ability of increasing concentrations of chimerophores cp9 (left) and its parent HDPs (pyrr_mod4_ middle, dc12_5_ right). Linearized pUC19 plasmid DNA (100 ng) was used.

### Chimerophores preserve the membrane translocation properties of their parent HDPs

Based on their favorable cytocompatibility, activity profiles, and non-lytic MOA, we selected cp6, cp9, cp13, cp15 and cp17 to investigate their membrane translocation mechanisms in *E. coli*. All these candidates contain at least one PrHDP, a peptide class typically exploiting an extended but very similar set of IM transporters^14,52,53^ (Extended Data Table 1). While cp6 and cp13 combine two PrHDP parents, potentially expanding uptake pathways via combination of IM transporter usage, cp9, cp15 and cp17 include a disordered non-PrHDP parent that bypasses IM transporters via self-promoted transient pore formation. We therefore investigated the effect of deleting *E. coli* IM transporters on chimerophore activity.

All *E. coli* knockouts lacked the primary PrHDP IM transporter *ΔsbmA* and one of the known secondary PrHDP transporters *ΔmdtM*, *ΔygdD* or *ΔyjiL* (Fig. 4c). In these mutants, the MIC of cp6 increased 4-fold, as observed for its PrHDP parents, onc_KR_ and api1B_G_ (by2-fold and >4-fold, respectively). As both parent modules utilize nearly identical IM transporters (Extended Data Table 1), this suggests that the chimerophore behaves consistently with the parent HDPs indicating no expansion of uptake pathways. Conversely, the MIC of the PrHDP-derived chimerophore cp13 remained unaffected, mirroring its parent bac7_PSL_ rather than its second parent, dro_WT_, whose MIC increased 6-fold (Extended Data Fig. 7f). This indicates that parental transport modalities can be inherited by chimerophores.

Interestingly, cp6_mod1_ showed high activity across all double knockouts, indicating reduced IM transporter dependence. Given that the Arg15/D-Arg and Arg19/D-Arg substitutions in its onc module have not been reported to enhance cellular uptake^39,46,47^, this effect is likely due to the additional cationic charge of the Orn residue in the api1B module, potentially increasing affinity for alternative IM transporters or direct membrane passage.

The activity of chimerophores cp9, cp15, and cp17 remained largely unaffected across knockouts (Fig. 4c; Extended Data Fig. 7g-i), in line with their design, which combines PrHDP parents relying on IM transporters with transporter-independent non-PrHDP parents. Specifically, the stable MIC of cp9 across all mutants mirrored the transporter-independent profile of its parent dc12_5_ (Fig. 4c), translocating via LPS binding and transient pore formation^54^, whereas its PrHDP parent pyrr_mod4_ showed an 8-fold MIC increase across all knockout strains. This suggests that cp9 effectively bypasses PrHDP resistance caused by IM transporter loss by exploiting the mechanisms of its non-PrHDP parent module. In cp9, the dc12_5_ parent may thus act as a vector enabling the uptake of the ribosomal inhibitor pyrr_mod4_ even if IM transporters are lost.

Unlike *E. coli*, *P. aeruginosa* exhibits high intrinsic resistance to PrHDPs thanks to a less permeable OM and the absence of SbmA. To evaluate if cp6 and cp9 retain a non-lytic MOA in this strain, we repeated the membrane damage assay using *P. aeruginosa* constitutively expressing mNeonGreen (mNG). cp6 and its parents remained non-lytic, suggesting that cp6 may have affinity for so far unidentified IM transporters in *P. aeruginosa*. Conversely, cp9 exhibited a membrane-damaging effect at the MIC (11% mNG+/PI-) (Extended Data Fig. 7e). As this lytic phenotype was not observed in *E. coli*, this suggests that the target bacterial cell envelope or its transport machinery determine the MOA of cp9. Importantly, despite the MOA shift, cp9 had low host-cell toxicity, overcoming a barrier otherwise common to lytic HDPs. Together, these results suggest that chimerization of non-lytic HDPs can reduce the reliance of PrHDPs on specific IM transporters by broadening their cellular uptake routes.

### Chimerophores retain the MOAs of parent HDPs

Next, we selected cp6 and cp9 to elucidate their molecular targets. While cp6 links a Type I and a Type II protein synthesis inhibitor (onc_KR_ and api1B_G_; PrHDP–PrHDP), cp9 was engineered from a Type I inhibitor (pyrr_mod4_) and a nucleic acid binding parent (dc12_5_; PrHDP–non-PrHDP fusion). We therefore assayed their affinity for these macromolecular targets in cell-free and live-cell models.

First, we assessed protein synthesis inhibition in an *E. coli in vitro* transcription/translation (IVTT) system using the firefly luciferase as a reporter (Fig. 4d). Both chimerophores showed a dose-dependent protein synthesis inhibition, with over 75% reduction in luminescence at 5 µM (Extended Data Fig. 8a). While the cp6 parent onc_KR_ efficiently inhibited luciferase synthesis (IC_50_=2.71 ± 0.61 µM), the second parent, api1B_G_, reached a non-dose-dependent saturation at 48% inhibition, preventing IC_50_ determination. This plateau is characteristic of Type II inhibitors, which permit residual luciferase synthesis prior to ribosome stalling^40,53,55–58^. Comparable to its Type I parent onc_KR_ (Fig. 4d), cp6 displayed an IC_50_ of 3.17 ± 0.45 µM) indicating inheritance of the parental inhibitory mechanism.

cp9 exhibited a 2.6-fold increase in potency (IC_50_ = 1.03 ± 0.05 µM) compared to its Type I inhibitor parent pyrr_mod4_ (IC_50_=2.67 ± 0.87 µM), alongside a stark increase in potency compared to its nucleic acid binding parent dc12_5_ (IC_50_ = 25.82 ± 8.90 µM). We attribute this marked improvement in translation inhibition partly to the modest baseline inhibitory value of dc12_5_, consistent with non-specific DNA-binding. Nevertheless, the finding that cp9 surpasses the potency of pyrr_mod4_ suggests an intramolecular synergistic interaction between the fused parent HDPs, which may simultaneously block transcription and translation by binding to both ribosomes and DNA.

Next, we evaluated the DNA-binding capacity of cp6 and cp9 in electrophoretic mobility shift assays (EMSA). Surprisingly, we observed complete DNA retardation at 4 μM for cp6 (Fig. 4e). Although neither parent was previously known to bind DNA^57^, onc_KR_ caused full DNA retardation at 8 μM whereas api1B_G_ showed no affinity. This suggests that cp6 inherited and doubled the DNA-binding activity of onc_KR_. Similarly, cp9 induced DNA retardation at 4 µM, indicating a 4-fold higher affinity than its DNA-binding parent dc12_5_ (Fig. 4f), while pyrr_mod4_ showed inactivity. Because both chimerophores utilize flexible linkers (GTG and GSG, respectively), this enhanced DNA affinity is not driven by a (poly)cationic linker sequence but instead points to an intramolecular synergy between the linked parent modules.

Collectively, these results demonstrate that both cp6 and cp9 inherit the ribosomal and DNA-binding mechanisms of their parents, supporting a multimodal MOA potentiated by intramolecular synergies.

To gain a broader understanding of its targets compared to its parents dc12_5_ and pyrr_mod4_, we next subjected cp9 to a proteome integral solubility alteration (PISA) assay (Extended Data Fig. 8b) using live cells for the identification of cellular pathways altered in response to peptide treatment, and cell lysates for the identification of putative direct binding events^59^. Gene set enrichment analysis (GSEA) of cp9-treated live cells (Extended Data Fig. 8b) revealed that ribosomal biogenesis and assembly pathways were destabilized by both cp9 and its ribosome-binding parent pyrr_mod4_, as indicated by negative normalized enrichment scores (NES). Furthermore, pathways involved in outer membrane (OM) structure and nucleic acid binding components were stabilized (positive NES) by both cp9 and dc12_5_, but not by pyrr_mod4_. Because the cellular uptake of dc12_5_ is facilitated by LPS binding, this provides evidence for the inheritance of this interaction by cp9. These results suggest that cp9 treatment shares a significant overlap in pathways perturbed by its individual parents, supporting retention of parental MOAs.

Finally, we analyzed the PISA profiles of cp9-treated cell lysates, revealing several proteins with significantly altered thermostability (Supplementary Table 4). Affected proteins included ribosomal components and modifying enzymes (namely, RpmF, RpsU, RlmD, and PrmB), supporting direct binding of cp9 to ribosomes. Additionally, cp9 showed interactions with nucleic acid binding (Hfq) and repair proteins (YoaA and RuvC), supporting cp9’s affinity to nucleic acids. Notably, cp9 showed interactions with membrane-associated proteins, including ATP-binding cassette (ABC) transporters and efflux pump components (YadG and CorC). Pyrr_mod4_ treatment induced thermal stability shifts in the cytosolic subunit of an ABC transporter (LptB), tRNA processing proteins (MiaA) and DNA repair proteins (Nfi), while dc12_5_ targeted gene expression regulators (Dcm, LeuO) and DNA repair factors (MutM, ElaB, ThiD, and MltC). Overall, these observations align with the pathways altered in live cells and illustrate a multimodal profile for cp9 encompassing the primary pathways of both parents.

### Chimerophores slow down resistance emergence in *P. aeruginosa*

To assess the propensity for evolutionary escape via genomic mutations, we performed 21-day serial passages using *P. aeruginosa* ATCC 27853 treated with either cp6 (onc_KR_–api1B_G_), cp9 (pyrr_mod4_–dc12_5_), or their parental peptides applied in equimolar ratios (Fig. 5a). As a positive control we used meropenem, known to rapidly induce resistance in *P. aeruginosa*. Each day, we determined the MIC and passaged the cultures growing at 0.5x the MIC into the next cycle. Resistance was defined as a MIC increase of at least 4-fold relative to day 1. By day 21, cultures exposed to cp6 and cp9 exhibited 4-fold and 1.3-fold MICs, respectively, indicating that resistance emerged for cp6 but not for cp9. The corresponding equimolar parent mixtures (additive controls) showed 1.3-fold and two-fold MIC increases. Although treatment with the parent mixtures did not drive the cells to resistance, the superiority of chimerophores is indicated by their 2- to 4-fold lower MICs. Conversely, meropenem induced resistance by day 3 and reached a 32-fold MIC increase by day 21 (Fig. 5a, b).

**Figure 5.**
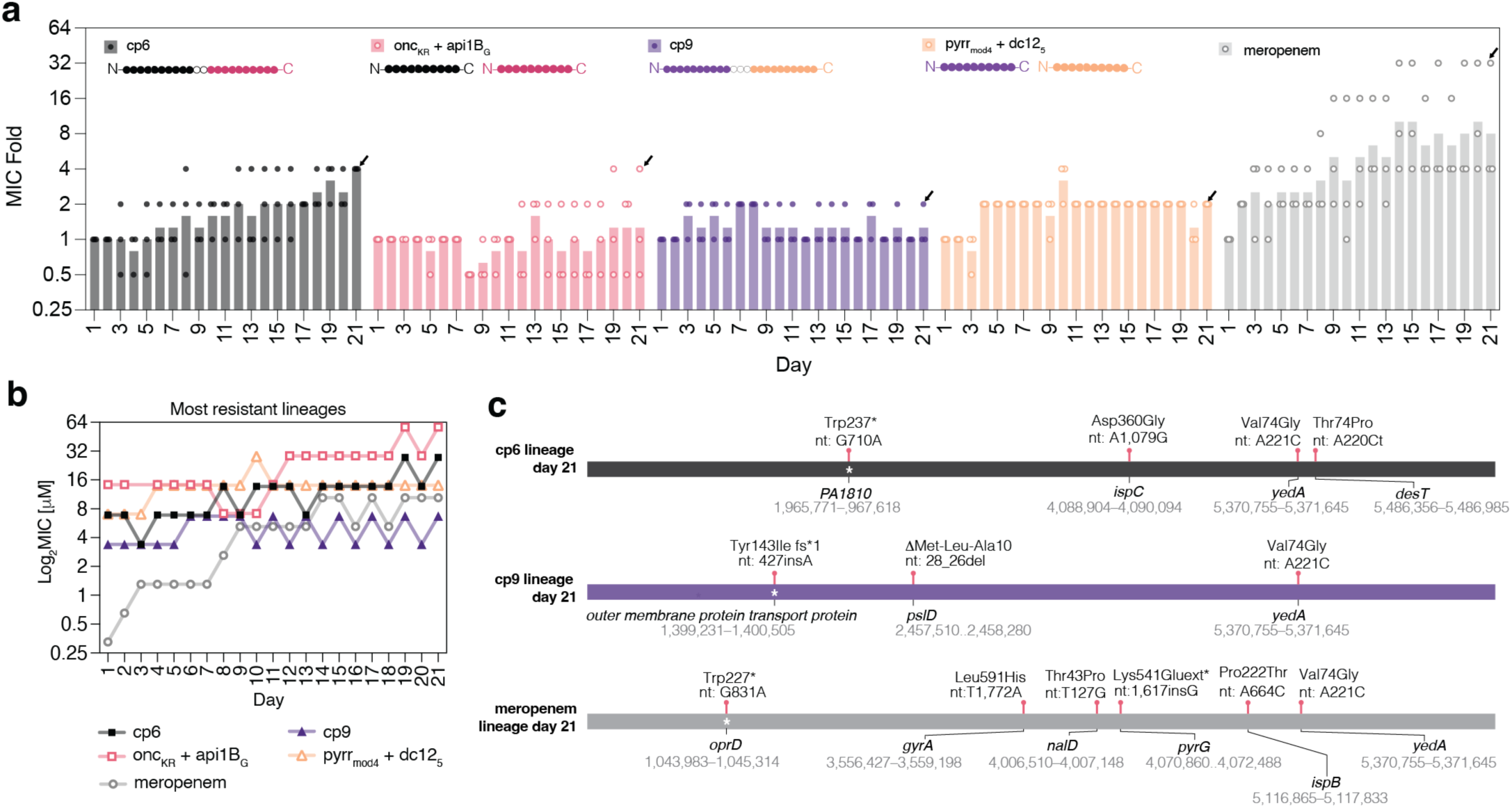
*P. aeruginosa* resistance during treatment with chimerophores and their parent HDPs applied at equimolar ratios. **(a)** Antibiotic resistance evolution profile of *P. aeruginosa* ATCC 27853 over 21 days. Bacterial lineages were evolved through a daily serial passage assay, tracking MIC increase over time. Each day, the subsequent MIC assay was inoculated using culture from 0.5xMIC well of the previous day. Resistance (defined as ≥4-fold MIC increase, dashed line) to chimerophores cp6 (black) and cp9 (purple) was compared to their respective parent HDP 1:1 molar mixtures (pink and orange), serving as a control for additive combination therapy, and the antibiotic meropenem (gray) as a positive control. MIC values are normalized to the MIC on day 1. Columns represent the mean of n=3 biological replicates, dots represent these replicates as independent lineages throughout the 21 days. Black arrows indicate lineages sequenced in (c). **(b)** MIC measurements over the 21 days of serial passage for the most resistant lineages derived from the treatments in (a), cp6 in black squares, parent combination onc_KR_ + api1B_G_ in open pink squares, cp9 in purple triangles, parent combination pyrr_mod4_ + dc12_5_ in open orange triangles and meropenem in open gray circles. **(c)** Genomic analysis of most resistant lineages following 21 days of serial passage from (b). Genome sequences from cp6 (black), cp9 (purple) and meropenem (gray) treatment were compared to the starting culture on day 1. Mutations are marked with red pins; white asterisks indicate premature stop codons; nt: nucleotide mutations; ins: insertion; del: deletion; fs*: frameshift leading to stop codon. Gene coordinates are based on *P. aeruginosa* PAO1 GenBank: GCA_000006765.1.

These results suggest that fusions of two PrHDPs with similar MOAs (e.g., onc–api1B in cp6) remain vulnerable to established resistance pathways, despite combining orthogonal Type I and Type II inhibitors. Furthermore, as pyrr_mod4_ lacked activity against *P. aeruginosa* (Fig. 3a), it likely exerts minimal selective pressure as a parent mixture. It is therefore plausible that the suppression of resistance observed with cp9 results from a multimodal MOA. By facilitating the transport of pyrr_mod4_, cp9 could bypass intrinsic resistance to pyrr_mod4_ and may enable concurrent targeting of both the ribosome and nucleic acids. Consequently, this multimodal MOA effectively maintains the potentiated antimicrobial activity and prevents the emergence of resistance.

Whole-genome sequencing of the most resistant lineage from each treatment group at day 21 (Fig. 5b, Supplementary Table 5) revealed that meropenem-resistant *P. aeruginosa* acquired mutations in the multidrug efflux pump regulator (NalD), the putative drug-efflux pump (YedA), and the essential DNA gyrase subunit A (GyrA). Additionally, we found a nonsense mutation in the OM porin OprD (Trp227*), well known to trigger carbapenem resistance^60^. Conversely, the lineages treated with parental mixtures only acquired a single missense mutation in YedA, suggesting that while mutations modulating efflux pumps accumulated, additional genomic alterations were restricted.

In comparison, cp6-treated lineage acquired multiple mutations. Specifically, in IspC, which is involved in isoprene-biosynthesis required for adjustment of the membrane fluidity, in the fatty acid synthesis regulator DesT, and in YedA, alongside a Trp237* nonsense mutation in the ABC transporter PA1810 (Fig. 5c). Thus suggesting that cp6’s evolutionary vulnerability does not stem from target-site mutations, but from its reliance on specific IM transporters, which is consistent with our observations in *E. coli* transporter knockouts. Conversely, cp9-treatment forced mutations in YedA and PslD, which are potentially involved in surface matrix modulation, alongside a nonsense mutation in the uncharacterized OM protein-transport protein PA1288. While these mutations may contribute to a minor increase in chimerophore tolerance, they failed to produce a resistance phenotype.

Taken together, the findings from MOA characterization to resistance assays indicate that cp9 exerts antimicrobial action against *P. aeruginosa* from multiple angles including membrane destabilization alongside binding of polynucleic acids and ribosomes. The combination of these effects imposes a distinct evolutionary pressure requiring multiple rather than single-locus mutations to acquire resistance. Of note, similar multi-target dynamics were reported for other HDPs such as buf2^61^ and LL-37^62^. This multi-targeted selective pressure may explain why evolutionary escape by the pathogen was prevented.

## Discussion

We demonstrate that a flexible chimeric design based on non-lytic HDPs, coupled with the Me^x^ high-throughput intracellular screening^26^ enables the rapid identification of multimodal chimerophores. The isolated variants combined (i) expanded activity spectra and enhanced potency; (ii) robust activity in serum alongside excellent therapeutic indices; (iii) dual translocation and dual intracellular targeting; and (iv) a significantly reduced propensity for the evolution of resistance. This library design proved particularly valuable for identifying PrHDP chimeras deemed as poor candidates due to blocked terminal amino acids essential for ribosomal exit tunnel penetration and binding. This highlights that rigid SAR paradigms may fail to predict chimerophore functionality. While some library members required specific linkers to rescue activity (e.g., api1B–dc12 chimerophores), others retained robust potency regardless of linker types despite blocked termini (e.g., api1B–bac5 chimerophores). Hence, constructing large, unbiased chimerophore libraries was essential to capture design trends and identify unexpectedly active variants.

Chimerophores comprising disordered non-PrHDPs (Arg-, Trp-, and Lys-rich) frequently failed to inhibit growth in the Me^x^ screen, particularly when placed at the N-terminus or if fused together. This likely reflects differential susceptibility to cytosolic peptidases; given that aminopeptidases are more abundant than carboxypeptidases in *E. coli*^63,64^, chimerophores with N-terminal disordered sequences likely have shorter half-lives, while those with dual disordered parents remain vulnerable from either end. Conversely, chimerophores with N-terminal or dual PrHDPs were more effective. The structural rigidity of PrHDPs, conferred by stable polyproline II helices^12,28,65,66^, likely contributes to resistance to aminopeptidases, extending stability in the Me^x^ screen. Furthermore, as PrHDPs target the ribosomal exit tunnel, they may gain additional protease protection via rapid post-translational target binding. Further studies on chimerophore metabolic stability and intracellular concentrations are necessary to determine whether these family-specific activity profiles stem from inherent Me^x^ screening limitations or represent genuine differences in intracellular target engagement.

Notably, linkers composed of basic residues were significantly overrepresented in the Me^x^-active fraction. As these linkers contribute +1 to +4 (e.g., GRG, RKRK) positive charges to the net charge of the chimerophore (with Me^x^-actives ranging from +2 to +22, Fig. 2i), they may enhance electrostatic interactions with polyanionic cytoplasmic components such as nucleic acids, ribosomes, and most globular proteins^67,68^. Such interactions could drive the formation of biomolecular condensates, ultimately triggering cell death as reported previously for LL-37^62^ and buf2^61^.

Alternatively, basic linkers (e.g., RKRK and HKHK) may be overrepresented because they serve as recognition sites for cytoplasmic endopeptidases like OpdB^28^, which protects cells from invading cationic peptides. While proteolysis is usually considered an inactivating resistance mechanism, biological precedent demonstrates that it can enhance peptide potency; for example, the pro-peptides bac5 and bac7 require extracellular cleavage to remove bulky C-terminal residues for activity potentiation^55,69^. This processing eliminates steric constraints on the N-terminal pharmacophore, yielding smaller fragments that can easily access the narrow ribosomal exit tunnel^34,70,71^. Similarly, we speculate that native bacterial peptidases may also cleave chimerophores at these basic linkers. Rather than inactivating the peptides, cleavage could liberate the individual parental modules, eliminating steric hindrance and allowing them to independently engage their respective intracellular targets.

This hypothesis is supported by our observation that certain chimerophores were active in Me^x^ only when connected by basic linkers containing potential OpdB cleavage sites rather than flexible linkers. For instance, api1B_WT_–bac5_258_ and api1B–ara1 combinations fuse a Type II PrHDP linked to the N-terminus of a Type I PrHDP, an arrangement expected to hinder ribosomal binding. Yet, their activity in Me^x^ suggests that intracellular cleavage at the OpdB linker site may unmask the previously blocked termini, enabling the liberated HDPs — either individually or synergistically—to bind the ribosome and inhibit translation. We propose that these early observations may inspire strategies to exploit strain-specific housekeeping proteases to potentiate chimerophore activity once they enter the cell.

By using non-lytic parent HDPs for library construction, we aimed to generate chimerophores that traverse the bacterial cell envelope using orthogonal mechanisms inherited from their parents. Because the Me^x^ selection workflow isolates intracellular activity rather than membrane translocation, it was initially unclear whether these chimerophores retained their cell penetrating ability if administered extracellularly. Strikingly, all 18 chimerophores that were chemically synthesized according to the template identified in the Me^x^ screen demonstrated potent *in vitro* antimicrobial activity when administered extracellularly. This 100% hit rate far exceeds those previously reported in other peptide self-screening assays^26,72^. Combined with the observation that 17 of these chimerophores retained a non-lytic MOA against *E. coli*, these results suggest that the modular incorporation of transport sequences in chimerophores successfully ensured their cellular uptake. This validates our chimerophore design as a reliable strategy for translating intracellular inhibitors into viable, membrane-permeable candidates.

Chimerophore activity profiles against the ESKAPE panel revealed a general trend toward enhanced potency compared to parent HDPs. In cases where one parent module was inactive against a specific strain, chimerophore activity matched (cp9) or closely approached (cp2) that of the active parent. Interestingly, several variants gained activity against strains outside of the combined parental range. For instance, cp14 gained activity against *K. pneumoniae*, despite both parents P-10 and indo being inactive. Similarly, seven chimerophores including cp6, cp15 and cp17 targeted *S. aureus*, despite originating from parents that primarily display activity against Gram-negative pathogens^16^. Crucially, this anti-Gram-positive activity could not be replicated using parent equimolar mixtures. We hypothesize this spectrum expansion results from the significantly higher net charge of chimerophores (e.g., +9 for cp6 in pH 7) compared to their individual parents (e.g., +7 for onc_KR_ and +2 for api1B_G_). The elevated cationic charge density likely enhances initial electrostatic interactions with surface LPS^54^ or LTA^67,73^, facilitating membrane-binding and self-promoted uptake^9,54^.

Alternatively, the high cationic density may trigger a membrane-disrupting MOA, as reported for other HDPs like bac7 against *P. aeruginosa*^19,37,74^ and for indo against *S. aureus*^75^. Indeed, we observed a distinct MOA shift in cp9 from non-lytic in *E. coli* to lytic in *P. aeruginosa.* In the latter organism, in the absence of PrHDP-required IM transporters, cp9 may have adopted and potentiated the transient pore formation mechanism of dc12_5_^54^ promoting a stronger pore formation in the IM of *P. aeruginosa*. We hypothesize that this MOA shift could also extend to other chimerophores with expanded activity spectra, suggesting that these fusions may adopt a lytic or a mixed mechanism in strains where intracellular targets are poorly accessible.

Crucially, chimerophores cp9, cp15, and cp17 robustly maintained activity in 20% serum, an environment known to compromise HDP activity through proteolytic degradation and a high ionic strength^5^. For cp9, this resilience likely stems from its parent, dc12_5_, whose high net charge and tryptophan content promote the interactions with OM components, counteracting the shielding effects of serum salts^54^. Cationic HDPs are often vulnerable to systemic serine proteases, typically cleaving at basic residues. Despite containing numerous predicted cleavage sites for serum serine proteases (e.g., thrombin and plasmin), these chimerophores showed stable activity suggesting significant resistance to such enzymes. This resistance may be mediated by the PrHDP parent pyrr_mod4_ in cp9, and the PRP-rich linkers in cp15 and cp17, exploiting Pro-rich motifs to sterically shield basic residues from protease active sites^12,76^. Thus, favorable intramolecular interactions and enhanced physicochemical properties can structurally protect chimerophores in physiological environments. Notably, because cp9, cp15, and cp17 consist entirely of canonical amino acids and unmodified termini, their stability may be further augmented using established modifications like D-amino acid substitution, N-terminal acetylation, or C-terminal amidation^39,77^.

Our observations further suggest that chimerophores exploit multimodal translocation mechanisms. For example, cp15 and cp17, both carrying the pharmacophore sequences of two PrHDP parents, maintained efficacy in *E. coli* lacking PrHDP-specific IM transporters^14,15,53^. The PRP motifs in their linkers likely contributed to this resilience, either by providing increased affinity to alternative unexamined IM transporters or enhancing membrane interactions for transporter-independent translocation^14,15^. By enabling alternative pathways for cellular entry like transient pore formation, chimerophores bypass transporter dependency, thereby reducing the likelihood of resistance via transporter loss. Future studies should elucidate these specific transport mechanisms across a broader range of Gram-negative and Gram-positive species.

EMSA and IVTT assays demonstrated that chimerophores cp6 and cp9 inherited the DNA-binding and translation-inhibiting mechanisms of the parental HDPs. Similarly, cp4, cp6 and cp13 involve two distinct binding sites within the ribosome exit tunnel likely occurring simultaneously, as they combine two PrHDPs targeting translation but via orthogonal translation mechanisms, Type I (translation elongation inhibition) and Type II (translation termination inhibition)^30,55,66,78–80^. Collectively, these bimodal MOAs likely contribute to the improved potency of many of the chimerophores synthesized in this study, including cp4, cp6, cp13, and cp15, as evidenced by substantially reduced MIC values against *P. aeruginosa* and *S. aureus* relative to their parents administered alone or in combination.

During 21-day serial passages, cp9 effectively suppressed resistance in *P. aeruginosa*. While tolerance to parental mixtures occurs via single mutations in a multidrug efflux pump^81,82^, chimerophore-treated lineages acquired mutations across multiple genes. In the cp6-resistant lineage, we identified mutations putatively modifying membrane fluidity and surface charge^83,84^, two established resistance mechanisms against cationic peptides^85^. Notably, truncation of the ABC-type IM transporter PA1810 suggests its involvement in cp6 cellular uptake, and its loss of function likely represents a primary driver of resistance to cp6. This aligns with the transporter-dependent activity of cp6 in *E. coli* and the well-documented reliance of PrHDPs on ABC-type IM transporters, like SbmA^14,15,52^. Conversely, *P. aeruginosa* remained susceptible to cp9 despite acquiring mutations in genes altering membrane permeability and cell surface composition, including porin truncation and altered LPS synthesis. This suggests that cp9 circumvents resistance mechanisms otherwise known to diminish the potency of not only cationic HDPs^85,86^ but also carbapenems^60^. This sustained potency likely arises from its multimodal MOA, exploiting a transporter-independent, pore-forming mechanism inherited from dc12_5,_ a potential second MOA from pyrr_mod4_, and a dual inhibition of essential intracellular processes.

Future work directed towards chimerophore development, using iterative design-test-learn-redesign cycles, may allow for the gradual optimization of these molecules toward further enhanced functional profiles. Furthermore, these high-quality datasets may provide valuable training sets for computational analyses and machine learning models. Ultimately, this approach supports the development of next-generation antibiotics capable of circumventing the early emergence of resistant pathogens typically associated with clinical use.

## Methods

### Bacterial strains

*E. coli* TOP10 was used as the Me^x^ screening strain. *E. coli* ATCC 25922 and *E. coli* BW25113 were used as references in MIC assays. MIC assays were done with several strains from the ESKAPE group: *Enterococcus faecium* LMG 16003*, Staphylococcus aureus* methicillin*-*susceptible ATCC 29213 *and S. aureus methicillin-*resistant ATCC 33591*, S. aureus* Triemli #104 (Triemli Hospital, Zurich), *Klebsiella pneumoniae* ATCC 13883*, Acinetobacter baumannii* carbapenem-resistant #105 (Kantonsspital St. Gallen)*, Pseudomonas aeruginosa* ATCC 27853*, P. aeruginosa* Triemli #082 (Triemli Hospital, Zurich), *Pseudomonas aeruginosa* meropenem-resistant 083M2 (Kantonsspital St. Gallen), *Enterobacter cloacae* E279 (Kantonsspital St. Gallen)*. E. coli* inner membrane knockouts were built using the KEIO collection strain *E. coli* JW0368 *ΔsbmA*^87^.

### Library design and cloning

Twelve ribosomally synthesized HDPs with established non-lytic MOAs and low cytotoxicity were selected as parents for chimerization. This set was supplemented with 44 engineered analogs identified in the literature for enhanced potency or designed in this study through the targeted substitution of potential OpdB cleavage sites with Gly residues (Supplementary Table 1). Chimerophore sequences were computationally designed in RStudio. Two principal chimeric architectures were developed, resulting in a library of 97,216 full-length and 2,016 PRP-linked chimerophores. Full-length chimerophores consisted of two parent HDPs tethered either directly or via one of 30 distinct linkers of 2 to 4 amino acids in length (Fig. 1b). PRP-linked chimerophores were designed by fusing the pharmacophore sequences of 13 parent HDPs while maintaining their native terminal orientations. Specifically, N-terminal pharmacophore sequences from bac7, bac5, onc, and P-10 were connected to C-terminal pharmacophore sequences from api1B, dro, buf2, dc12, indo, PR-39 and pyrr (Extended Data Fig. 1a). These modules were connected by one of 48 distinct 11-residue PRP-linkers. These linkers were derived from a consensus motif identified via multiple sequence alignment of the complementary sequences of the 13 parents, using the T-Coffee algorithm in JalView (Extended Data Fig. 1b).

#### Sequence preparation and oligo synthesis

Amino acid sequences of chimerophores were back-translated and codon optimized for *E. coli* K12 in Geneious Prime, while excluding internal restriction sites for *HindIII* and *PstI*. Coding sequences were flanked *in silico* by universal Me^x^ primer binding sites^26^ containing *PstI* (Primer 1) and *HindIII* (Primer 2) recognition sites to facilitate downstream cloning (Supplementary Table 6). The resulting primer-flanked chimerophore coding library was synthesized as a single-stranded DNA (ssDNA) oligo pool by TWIST Biosciences.

#### Molecular cloning and library construction

The ssDNA oligo pool was amplified by PCR using Phusion High-Fidelity PCR master mix with GC buffer (New England Biolabs, M0532L). Reactions contained 10 µM of Me^x^ Primers 1 and 2 (Supplementary Table 6) and 20 ng of template ssDNA in a 50 µL volume. The resulting double-stranded DNA was purified using QIAquick gel extraction kit (Qiagen, 28704) and concentrated with the DNA clean & concentrator-5 (Zymo, D4004). For ligation, 500 ng of PCR-amplified library and 1 µg of the pBAD expression vector were subjected to double digestion with *PstI*-HF (New England Biolabs, R3140S) and *HindIII*-HF (New England Biolabs, R3104S) according to the manufacturer’s protocol. Digested fragments were purified as described for PCR products and ligated at a 7:1 (insert:vector) molar ratio with T4 DNA ligase (New England Biolabs, M0202S) for 14 h at 16°C. The ligation product (800 ng) was used to transform 200 µL of electrocompetent *E. coli* TOP10 cells (ThermoFisher, USA) via electroporation (Bio-Rad Micropulser). Recovered cells were plated on LB agar supplemented with 100 µg/mL carbenicillin and 0.2% (w/v) glucose, and incubated overnight at 30°C. Colonies were harvested as a pool, adjusted to an OD_600_ of 1 in LB medium containing 20% (v/v) glycerol, and stored as 1 mL aliquots at -80°C.

### Me^x^ growth assay

To assess the antimicrobial activity of the peptides in the library, we performed massively parallelized growth assays (Me^x^) as previously described^26^. In short, Me^x^ allows for recording of growth curves of each of our 97,216 peptide-expressing *E. coli* strains and the identification of growth-inhibiting peptides by next-generation sequencing (NGS). For this, we inoculated liquid cultures with approximately 2.5 billion transformed cells in triplicates and induced peptide synthesis in the early exponential growth phase (OD_600_ ∼ 0.08; Extended Data Fig. 2a). We monitored the growth of the cultures for 4.5 h post-induction, while harvesting bacterial samples every 1.5 h for NGS analysis. Following sample collection, plasmid DNA was extracted, and Illumina barcodes were added by PCR. Finally, equimolar amounts of each sample were pooled for NGS.

### NGS and data analysis

The plasmid samples were sequenced on an Illumina NovaSeq platform (Functional Genomics Center Zurich) using paired-end sequencing (2 x 150 bp), generating 200 million reads. We chose this read length to ensure complete coverage of the 194 bp peptide-encoding sequences. For data processing, we merged the paired-end reads using PEAR^88^. The resulting full-length reads were mapped to our peptide library reference database using BBMap^89^. To ensure high-quality data, we removed ambiguous mapping reads. Read counts for each peptide were combined into a single table and assigned a unique identifier (ID). Only peptides with a minimum of 15 reads at the initial time point (t_0_) were used for analysis (Extended Data Fig. 2b). The differential abundance of peptide-encoding DNA (at t_3_ vs. t_0_) was analyzed in RStudio (R version 4.4.3) using the DESeq2 package^90^, following a previously described method^26^. Growth-inhibiting activity was defined as an LFC below –1 with an adjusted p-value of less than 0.05 (one-tailed Wald’s test). To quantify the time-dependent loss of plasmid-carrying culture, the estimated OD_600_ of the pooled culture (OD_ID_), calculated as the product of the OD*_t_* and the LFC*_t_* at each time point, was used to generate growth curves of each variant.

Chimerophores were grouped into activity categories to distinguish between highly active, moderately active, and inactive chimerophores based on their growth-inhibitory activity while considering key physicochemical properties that may influence their synthesis and clinical potential. Active chimerophores were categorized into categories 1 to 4, with decreasing activity: category 1 (LFC < -6), category 2 (-6 < LFC < -4), category 3 (-4 < LFC < -2), and category 4 (-2 < LFC < -1). Inactive chimerophores were placed in categories 5 (-1 < LFC < 0) and 6 (LFC > 0).

A two-sided Fisher’s exact test was applied to assess the enrichment profile of parent HDPs in different configurations as well as of linker types in the group of Me^x^ actives, as previously described^26^. The null hypothesis was evaluated at a confidence level of 95%, and adjustment for multiple testing was performed using the Benjamini-Hochberg method. Parent HDP combinations and linker types with adjusted *p* < 0.05 were considered significantly enriched (over- or under-represented) in the Me^x^ active set compared to the entire library.

### Selection of chimerophores for chemical synthesis

For *in vitro* characterization, a representative subset of 18 chimerophores was selected, prioritizing physicochemical thresholds for optimized solid-phase peptide synthesis (SPPS) and *in vitro* handling. Physicochemical properties of the whole chimerophore library were predicted in RStudio using the Peptides package^43^. To mitigate peptide aggregation and facilitate purification via reverse-phase high-performance liquid chromatography (RP-HPLC), candidates were restricted to a net charge below +11 (calculated at pH 7 using the EMBOSS pK scale) and a negative Grand Average of Hydropathicity (GRAVY) score (Kyte-Doolittle scale). Sequence length was capped at 50 amino acids to ensure feasibility for SPPS^42^. Additionally, an instability index below 41 was applied to prioritize candidates with high predicted *in vitro* stability and potentially extended *in vivo* half-lives^44^. Selected chimerophores and their parent HDPs were chemically synthesized by Wuxi AppTec (China) with a purity of >95% and provided as trifluoroacetate salts.

### Monoseptic growth assays

We validated the growth-inhibiting activity of chimerophores of Me^x^-positive strains by using four randomly selected *E. coli* TOP10 strains carrying a peptide per activity category 1 to 5 and grown in separate wells of a microtiter plate. Peptide intracellular synthesis was induced at OD_600_ 0.08 with 0.3% (v/v) L-arabinose and incubated at 37°C in a plate reader (Tecan M1000) with orbital shaking at 1.5 mm amplitude, and absorbance measurements were made every 1000 s, for 24 h. The acquired data was trimmed at 6 h for comparison to Me^x^ and then analyzed using GraphPad Prism 10.

### MIC determination using chemically synthesized peptides

We determined the minimal inhibitory concentration (MIC) for the selected HDPs as previously described^91^. Briefly, 5 mL bacterial cultures were grown at 37°C for 16 h in 15 mL culture tubes in a shaker (250 rpm). Two-fold dilution series of HDPs in 25% Mueller Hinton Broth were prepared in 96-well plates (Thermo Scientific, 243656) using an Opentrons OT-2 robot. Bacterial cultures grown overnight were diluted to 1x10^6^ CFU/mL and 50 µL were added to each well for a final volume of 100 µL per well, and a starting concentration of 5x10^5^ CFU/mL. The plates were incubated at 37°C for 20 h. The absorbance at 600 nm was measured in a plate reader (Tecan Spark), and the MIC of the HDPs was determined as the concentration where the OD_600_ was at least 95% lower than that of untreated cells.

### *In vitro* cytotoxicity assay

The cytotoxicity of selected chimerophores in human embryonic kidney cells was assessed as previously described^92^. In brief, HEK293T cells (Merck, 12022001) were cultivated in Dulbecco’s Modified Eagle Medium (DMEM), high glucose (Gibco, 11965092) supplemented with 10% fetal bovine serum (Merck, F2442). Before seeding, cells were washed once with one volume of Dulbecco’s Phosphate-Buffered Saline (DPBS) (Merck, D8537) and resuspended in prewarmed DMEM. For the cell viability assay, 25,000 HEK293T cells were seeded into 96-well plates (Merck, CSL3599) with 100 µL DMEM. The last row of each plate contained DMEM only. The cells were incubated for 24 h at 37°C with 5% CO_2_ before treatment with HDPs. Two-fold dilutions of HDPs were prepared in 50 µL in v-bottom 96-well plates (Thermo Scientific Nunc, 249935) as described above for MIC assays, except 50% DMSO was added to column 11 as positive control for toxicity. To start the assay, 50 µL of cells were added to each well in columns 1-11. The plates were incubated for 24 h at 37°C and 5% CO_2_.

To determine the viability of HEK293 cells treated with HDPs, the Cell Proliferation Kit I MTT (Roche, 11465007001) was used following the manufacturer’s instructions. Briefly, MTT (3-(4,5-dimethylthiazol-2-yl)-2,5-diphenyltetrazolium bromide) was filtered through a 0.2 µm acetate filter, 10 µL were added to the cells and the plates were incubated for 4 h at 37°C and 5% CO_2_ in a non-shaking incubator (Eppendorf, CellXpert). Afterwards, 100 µL of solubilization buffer was added to the cells and the plates were incubated for another 24 h at 37°C and 5% CO_2_. The absorbance at 575 nm was measured in a plate reader (Tecan Spark) with a reference wavelength of 690 nm. The IC_50_ was determined by nonlinear regression of the dose-response curves in GraphPad Prism 10.

### Hemolysis assay

Hemolysis assays were performed as described previously^93^ with some modifications. Human fresh blood samples (Blutspende SRK beider Basel) from three different donors were collected in tubes containing ethylenediaminetetraacetic acid (EDTA) as anticoagulant. Samples were centrifuged at 1,500 × g for 5 min and the resulting plasma fraction was removed from the samples. The pellets containing red blood cells were washed 3 times with an equal volume (200 µL) of PBS (Gibco, 10010023). The washed red blood cells were diluted with 1x PBS to make a 2% suspension. The assay was performed in 384-well microplates with cover (Greiner Cat. No. 781801) in a final volume of 50 µL. The HDPs were tested in serial 1:1 dilution in PBS, with final concentrations ranging from 128 to 0.5 µM. Melittin was used as a reference for a hemolytic peptide and 1% TritonX100 (Sigma-Aldrich, 9036-19-5) and PBS were used as positive and negative (diluent) controls, respectively. The plates were incubated for one hour at 37°C. Finally, the plates were centrifuged at 1,500× g for 5 min at room temperature (Eppendorf, Centrifuge 5920 R). Supernatants were removed from all plate wells and transferred to a new plate for absorbance measurements. The plate was centrifuged at 1,000 ×*g* for 1 min to remove air bubbles and optical density was measured at 405 nm using a plate reader (Tecan Spark). The percentage of hemolysis was calculated as:

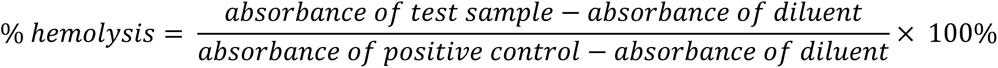

### MIC assays in human serum

The stability of selected chimerophores was assessed by MIC assays using *P. aeruginosa* ATCC 27853 and *S. aureus* ATCC 29213 in the presence of 20% human serum. Serum from three donors was obtained from Blutspende SRK beider Basel (Switzerland). The serum was defrosted at 37°C and then centrifuged at 15,871 ×*g*, 4°C for 10 min (Eppendorf, Centrifuge 5424 R) to remove lipids. The lipid-free supernatant was collected and incubated at 37°C for 15 min. Peptide dilution series were prepared in triplicates in 96-well plates (Thermo Scientific, 243656) as described for MIC assays (see above). Bacterial cultures were grown overnight at 37°C and then diluted to 1x10^6^ CFU/mL in a 40% serum solution in 25% MHB. The assay was initiated upon the addition of 50 µL of diluted bacterial cultures to 50 µL of peptide dilution series. The plates were incubated for 20-24 h at 37°C and the absorbance at 600 nm was measured in a plate reader (Tecan Spark).

### Membrane damage assay and flow cytometry data analysis

The membrane-damaging effect of parent pharmacophores and chimerophores was assessed as described previously^26^ with some modifications. Briefly, *E. coli* TOP10 [pSEVA271-GFP] and the peptide two-fold dilutions were prepared in duplicates as described for the MIC assays starting with a concentration range of 16xMIC to 0.03xMIC. The bacterial culture was diluted to 1x10^6^ CFU/mL in 25% MHB containing 1 µg/mL propidium iodide (PI) and added to the peptide dilution series. The assays were done in a 0 of 100 µL, the plates were incubated at room temperature for 1 h. The uptake of PI and release of GFP in 10,000 single cells per well was assessed by flow cytometry using a Fortessa Analyzer equipped with a 488 nm laser with 530/30 nm bandpass filter and a 579 nm laser with 610/20 nm bandpass filter (BD LSR Fortessa SORP, BD Biosciences). The fractions of PI-positive/negative and GFP-positive/negative were determined using FlowJo V10 (BD Biosciences).

### Construction and use of knockout *E. coli* strains

Inner membrane double knockouts *E. coli* BW25113 *ΔsbmA ΔygdD*, *E. coli* BW25113 *ΔsbmA ΔmdtM* and *E. coli* BW25113 *ΔsbmA ΔyjiL* were created via homologous recombination as previously described^94^. Briefly, *E. coli* BW25113 *ΔsbmA* (JW0368) was transformed with helper plasmid pKD46, which carries the required lambda red genes, and incubated at 30°C. The chloramphenicol resistance gene flanked by flip-recombinase target (FRT) sites and 50 bp homologous to upstream and downstream regions of the genes of interest were purchased as gene fragments (TWIST Biosciences, USA). Expression of pKD46 genes was induced with 0.3% (v/v) L-arabinose at OD_600_ 0.6, the cells were then made electrocompetent and 50 µL cell aliquots were electroporated with 500 ng of one of the three chloramphenicol gene fragments. Transformants were selected on LB plates with 50 µg/mL kanamycin and 34 µg/mL chloramphenicol and re-streaked twice before confirmation by colony PCR using the Phusion High-Fidelity PCR master mix with GC buffer (New England Biolabs, M0532L) and the primers mdtM-fw, mdtM-rev, ygdD-fw, ygdD-rev, yjiL-fw and yjiL-rev (Supplementary Table 6). Positive clones were then confirmed by whole-genome sequencing BacterialSeq (Microsynth, Switzerland) using genomic DNA purified using the Quick DNA Fungal/Bacterial miniprep kit (Zymo, D6005). Confirmed clones were incubated at 42°C to lose the pKD46 plasmid.

### Inhibition of *in vitro* translation assay

The *in vitro* translation reactions were performed as described before^56^ using the PURExpress system (New England Biolabs, E6800L). A plasmid pET28a(+) encoding the Firefly luciferase (Fluc) gene was used as template DNA. Each 5 µL reaction contained 10 ng of template DNA and the PURExpress reaction mix. Then, 1 µL of peptide was added to final concentrations of 0, 5, 10, 30, 60, 150 µM. Reactions were incubated for 30 min at 32°C. To stop the reactions, 2 µL of each reaction mixture was transferred to 8 µL of kanamycin (50 mg/ml) in a white 96-well plate. Finally, 40 µL of SteadyGlo luciferase substrate (Promega, E2510) was added to each well. Luminescence was measured in a plate reader (Tecan Spark).

### Electrophoretic mobility shift assay (EMSA)

The DNA-binding activity of chimerophores and their parents was assessed in an electrophoretic mobility shift assay as described before^9,54,56^. In brief, *E. coli* TOP10 carrying a pUC19 plasmid vector (Thermo Scientific) was cultivated overnight at 37°C. Plasmid DNA was extracted using a Zyppy plasmid miniprep kit (ZYMO, D4036)^5,19,20^.

Increasing concentrations of peptides (0, 2, 4, 8, 16, 32 µM) were incubated at 37°C for 1 h with 100 ng of pUC19 plasmid DNA (60.4 fmol) linearized with HindIII-HF (NEB, R3104S). All reactions were performed in 15 µL with binding buffer containing 10 mM Tris-HCl, 5% glycerol, 50 μg/mL BSA, 1 mM EDTA, and 20 mM KCl pH 8.0. The SDS-free orange DNA dye (Thermo Scientific, R0631) was used for sample visualization. Retardation of DNA due to HDP binding was assessed by gel electrophoresis in a 1% agarose gel in 0.5x TAE buffer, run at 60 V for 90 min, and then imaged using a FastGene FAS-Digi PRO system (NIPPON Genetics). All images were processed using ImageJ 1.54g.

### Analysis of proteome integral solubility alteration (PISA)

PISA was performed as described previously^59,95^. *E. coli* ATCC 25922 was grown in MHB at 37°C to an OD_600_ of 0.6. For PISA with cells, 1.5 x 10^9^ cells were incubated with 0, 4, 16, or 64 µg/mL HDP or chimerophore in 200 µL 25% MHB for 90 min at 37°C. For PISA with cell lysates, 7.5 x 10^9^ cells were resuspended in 200 µL lysis buffer (25 µg/mL lysozyme, 1x cOmplete EDTA-free protease inhibitors (Roche, 11836170001), 250 U/mL benzonase (Millipore, E1014), 1 mM MgCl_2_, in PBS) and incubated at 25°C for 20 min. Lysates were frozen in liquid nitrogen and then thawed at 25°C for three cycles. Cell debris was removed by centrifugation at 10,000 ×*g* for 5 min at 4°C (Eppendorf, Centrifuge 5424 R). Supernatants were incubated with 0, 4, 16, or 64 µg/mL parent HDP or chimerophore for 30 min at 37°C.

For thermal gradient treatment, samples (cell and lysate) were aliquoted row-wise into 96-well LoBind PCR plates (Eppendorf, 0030129504) from column 2 to 11. The plates were exposed to a thermal gradient (51-62°C) for 3 min in a qPCR machine (qTOWERiris) followed by 3 min at 25°C and then frozen in liquid nitrogen. The samples were thawed on ice, pooled and ultracentrifuged at 100,000 x g for 20 min at 4°C. Protein concentration was measured, and 10-50 µg of total protein was diluted in 70-100 µL SDS lysis buffer (5% SDS, 10 mM TCEP, 100 mM triethylammonium bicarbonate–TEAB) for protein digestion.

#### Protein digestion

Samples in SDS buffer were heated at 95°C for 10 minutes and then alkylated with 1 µL 1 M iodoacetamide at 25°C for 30 minutes (500 rpm). Each 25 µL of sample was acidified with 2.5 µL of 12% phosphoric acid (1.2% final concentration). Then, S-trap buffer (90% methanol, 100 mM TEAB pH 7.1) was added in a 6:1 ratio and vortexed. The samples were loaded onto an S-trap micro column (Protifi), centrifuged at 4,000 ×*g* for 1 minute, and the flow-through was removed. Samples were washed three times with 150 μL S-trap buffer, centrifuging at 4,000 ×*g* for 1 minute after each wash (Eppendorf, Centrifuge 5424 R). For digestion, 20 μL of digestion buffer (50 mM TEAB pH 8.0) with 1 µg Trypsin (Thermo Scientific, 90058) was added to the columns and incubated for 1 hour at 47°C in the dark. Digested peptides were eluted sequentially with 40 μL digestion buffer, 40 μL of 0.2% formic acid and 35 μL of 50% acetonitrile with 0.2% formic acid. Centrifugation steps were done at 4,000 ×*g* for 1 min (Eppendorf, Centrifuge 5424 R). The samples were dried by speed vac. Peptides were dissolved in LC buffer A, ultrasonicated briefly (10 seconds, 1400 rpm, 25°C), and concentration adjusted to 0.5 μg/μL. Samples were analyzed by mass spectrometry-based proteomics as previously described^59,95^.

#### Proteomics data analysis

Data processing began by selecting identified proteins that passed the Proteomics Core Facility (PCF) quality filters (Supplementary Fig. 5). To ensure high-confidence quantification, we restricted the analysis to proteins with a unique peptide count (UPC) >2. Differential abundance and solubility changes were determined using an adjusted p-value <0.05 and a log_2_(fold change) (LFC) threshold of ±0.5. Proteins exhibiting infinite LFC values, representing instances where a protein was uniquely identified in only one experimental condition, were included in the analysis only if they were supported by a UPC >2. To elucidate the biological pathways affected by chimerophore treatment, Gene-Set Enrichment Analysis (GSEA) was performed on selected conditions. These analyses were conducted using the clusterProfiler^96^ version 4.14.6 using RStudio.

### Resistance studies

Resistance evolution was assessed in *P. aeruginosa* ATCC 27853 using a previously described MIC assay format^97^. Briefly, an initial MIC was determined for each antimicrobial peptide in 25% MHB. Subsequently, each day for 21 days, cultures from the highest concentration that supported growth (0.5× MIC) were diluted 10,000-fold and used to inoculate a fresh two-fold dilution plate of the respective peptides. After 22 hours of incubation, absorbance at 600 nm was measured using a Tecan Spark plate reader. Meropenem served as a reference compound. On day 21, cells from 0.5x MIC wells were collected, supplemented with 20% glycerol and stored at -80°C. The most resistant lineage from each HDP treatment and the original wild-type strain were grown in 3 mL of LB medium at 37°C in a shaking platform at 250 rpm (Rotamax 120, Heidolph) to an OD_600_ of 1. Genomic DNA was extracted using a GenElute Bacterial Genomic DNA Kit (Sigma, NA2110). Specifically, 2.5 mL cultures were used, and the final elution was performed with 100 μL of sterile distilled water. The genomic DNA was then sent to BacterialSeq (Microsynth, Switzerland) for whole-genome sequencing. To identify potential resistance-associated mutations, whole-genome sequences were aligned using Mauve Alignment^98^ in Geneious Prime 2025.1.2.

## Acknowledgements

We acknowledge Next-Generation Sequencing support by Dr. Maria Domenica Moccia and Dr. Natalia Zajac (Functional Genomics Center Zurich, Zurich, Switzerland). We thank Dr. Qun Ren Zulian (EMPA, St. Gallen, Switzerland) for kindly providing *P. aeruginosa* and *S. aureus* clinical isolates, as well as scientific advice. We thank Leonard Fröhlich for his assistance during activity assays with parent equimolar mixtures and for providing the PAO1 mNeonGreen strain. We thank Dr. Danilo Ritz and Dr. Alexander Schmidt (Proteomics Core Facility, Biozentrum, Basel, Switzerland) for excellent support in Proteomics experiments. Schemes in Figure 1, Figure 4a, and Extended Data Figure 7f were created with Biorender.com and edited in Adobe Illustrator.

## Author contributions

L.A.E. and M.H. conceptualized the study and wrote the manuscript. L.A.E. performed all experiments, analyzed the data. S.L. performed cytotoxicity tests in HEK 293 cells, and L.A.E. analyzed the data. M.H. and S.P. supervised experiments. S.P. provided funding. All authors revised the manuscript.

## Extended data

**Extended Data Table 1.**
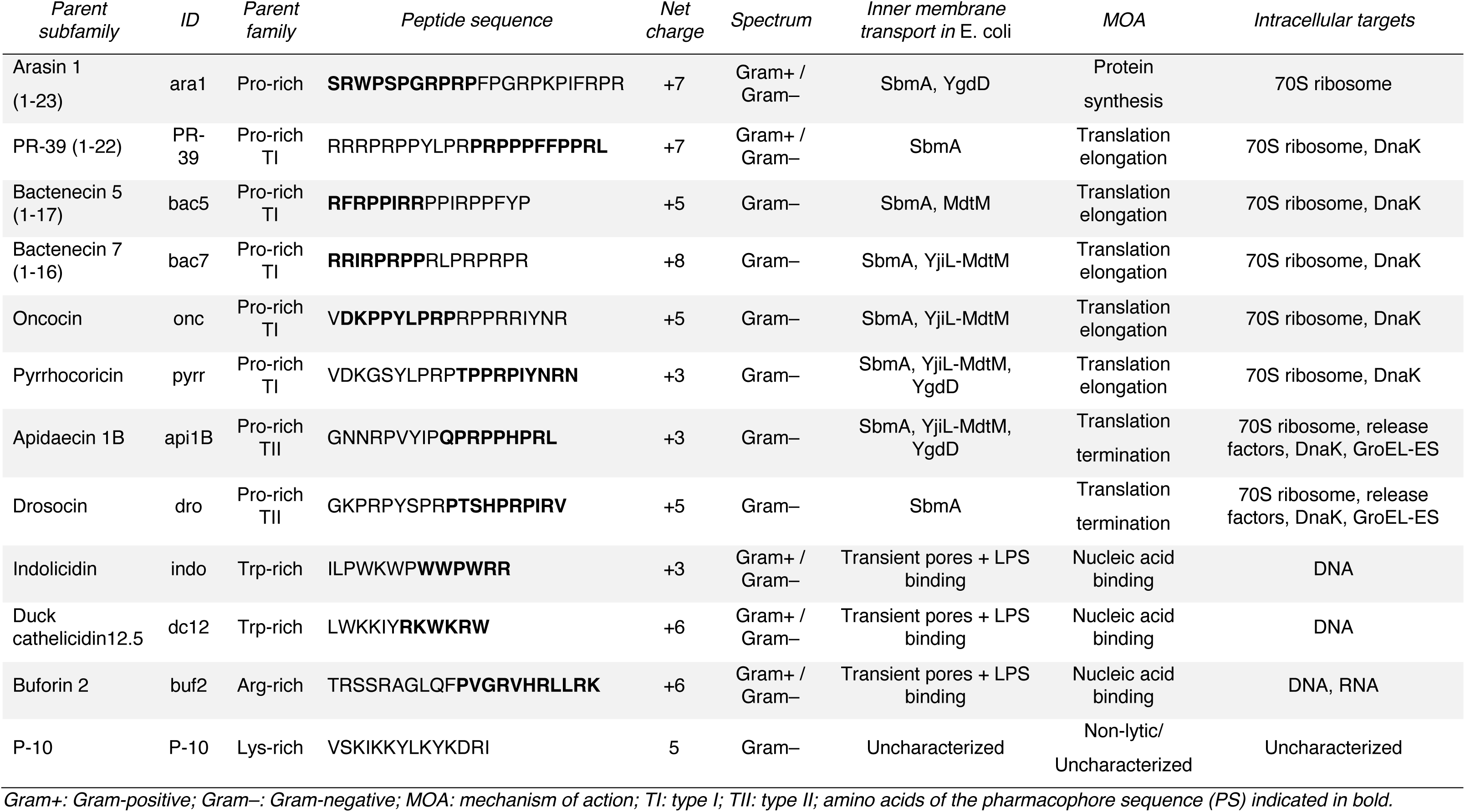
Parent HDP subfamily information.

**Extended Data Table 2.**
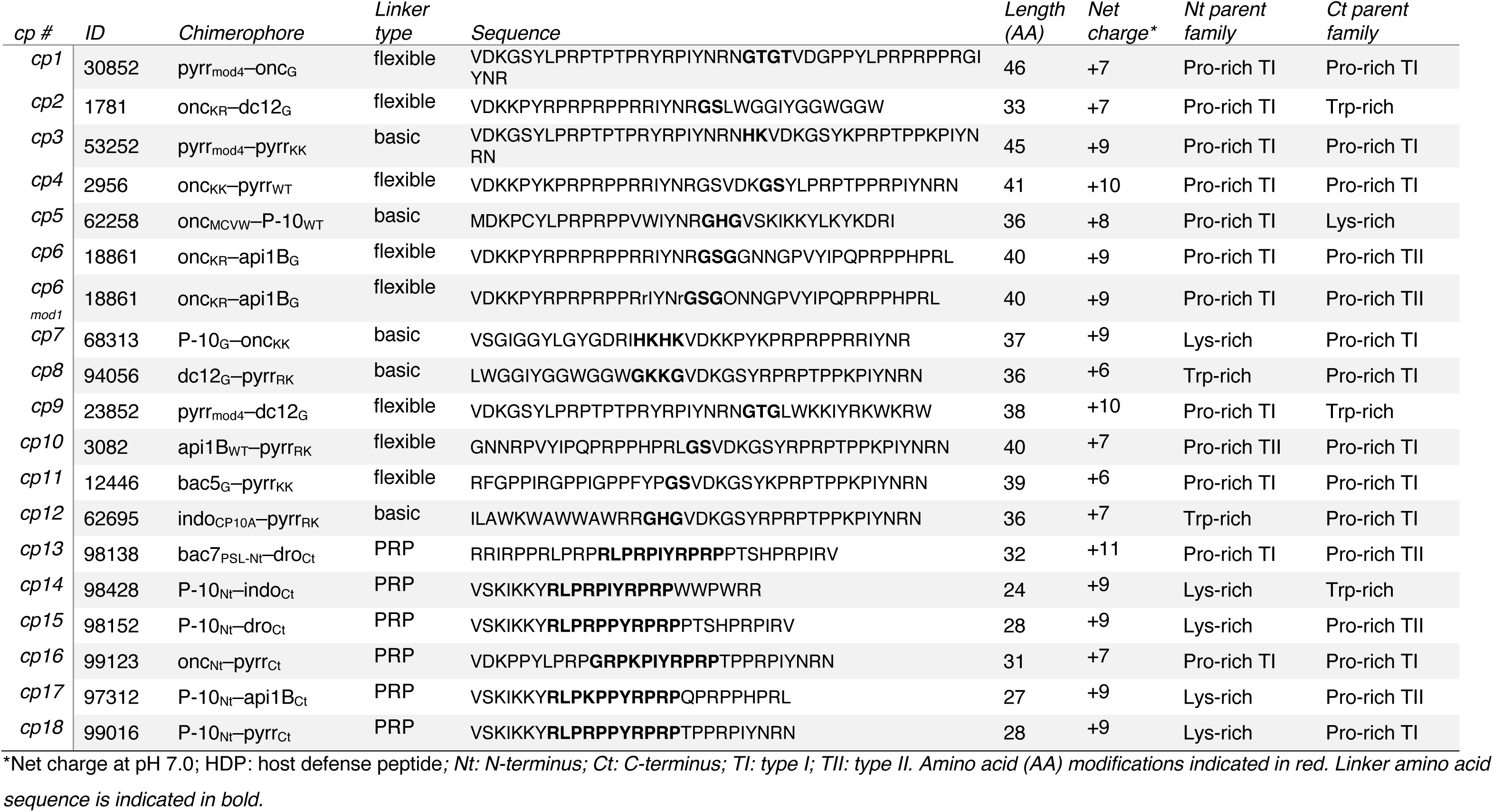
Information on 18 selected chimerophores.

**Extended Data Figure 1.**
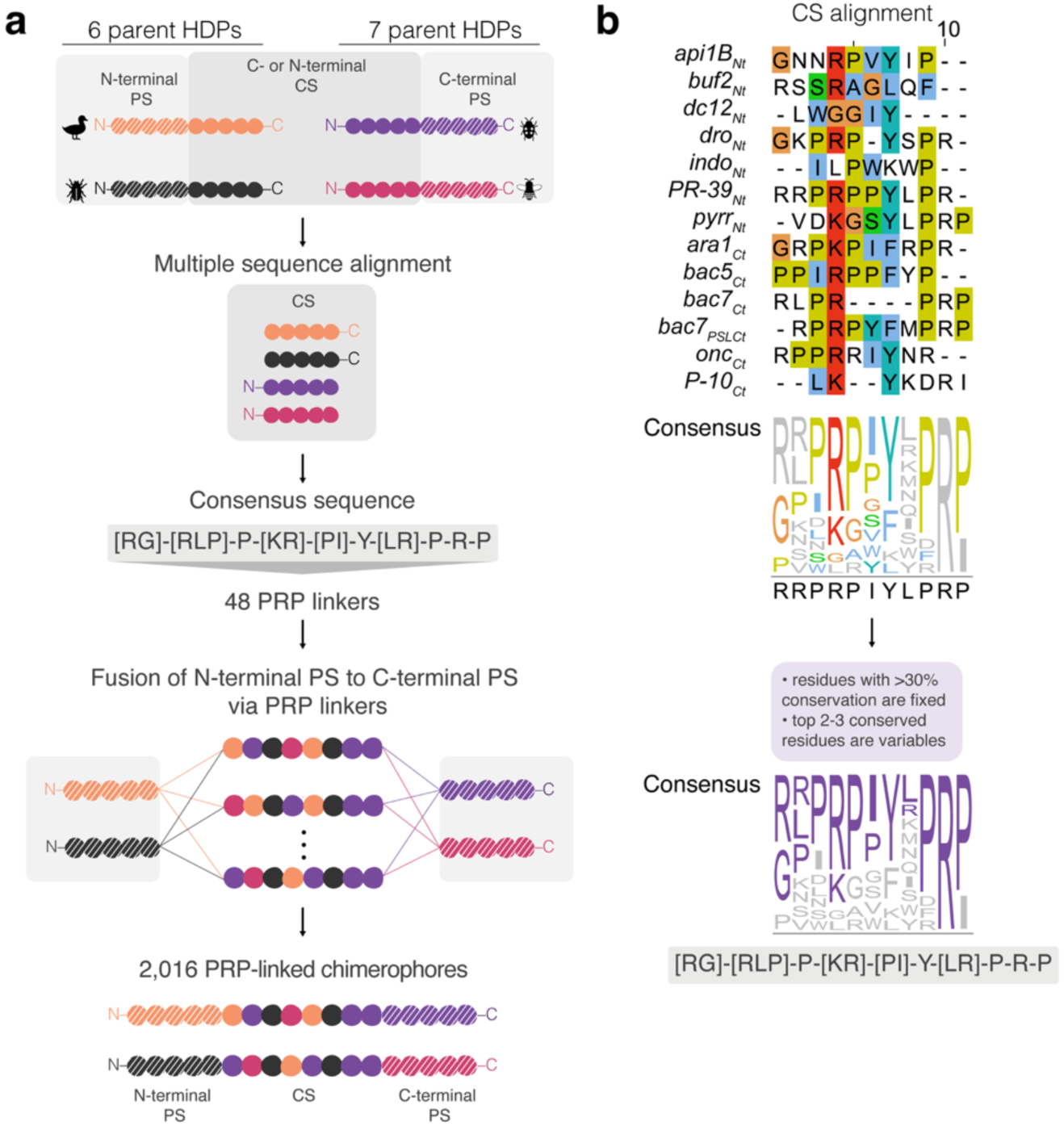
Design of PRP-linked chimerophores. **(a)** Combinatorial assembly of the PRP-linked library. The complementary sequences (CS) of 13 parent HDPs were aligned to derive an 11mer consensus sequence, which was used to produce 48 unique PRP linkers. These linkers were then used to bridge the pharmacophore sequences (PS) of the 13 parents (6 N-terminal PS: ara1_WT_, bac5_WT_, bac7_WT_, bac7_PSL,_ onc_WT_, and P-10_WT_; and 7 C-terminal PS: api1B_WT_, buf2_WT,_ dc12_5_, dro_WT_, indo_WT_, PR-39_WT_, pyrr_WT_) resulting in 2,016 PRP-linked chimerophores. In these tripartite constructs, the native orientation of the PS was preserved: N-terminal PS remained at the N-terminus, and the C-terminal PS remained at the C-terminus. **(b)** Sequence alignment and consensus derivation. Clustal Omega alignment of CS sequences extracted in (a); residues are colored by Clustal properties. The 11mer consensus (purple) was defined by fixing positions with >30% conservation and allowing the remaining positions to vary among the 2-3 most frequent residues, yielding 48 possible PRP linker sequences.

**Extended Data Figure 2.**
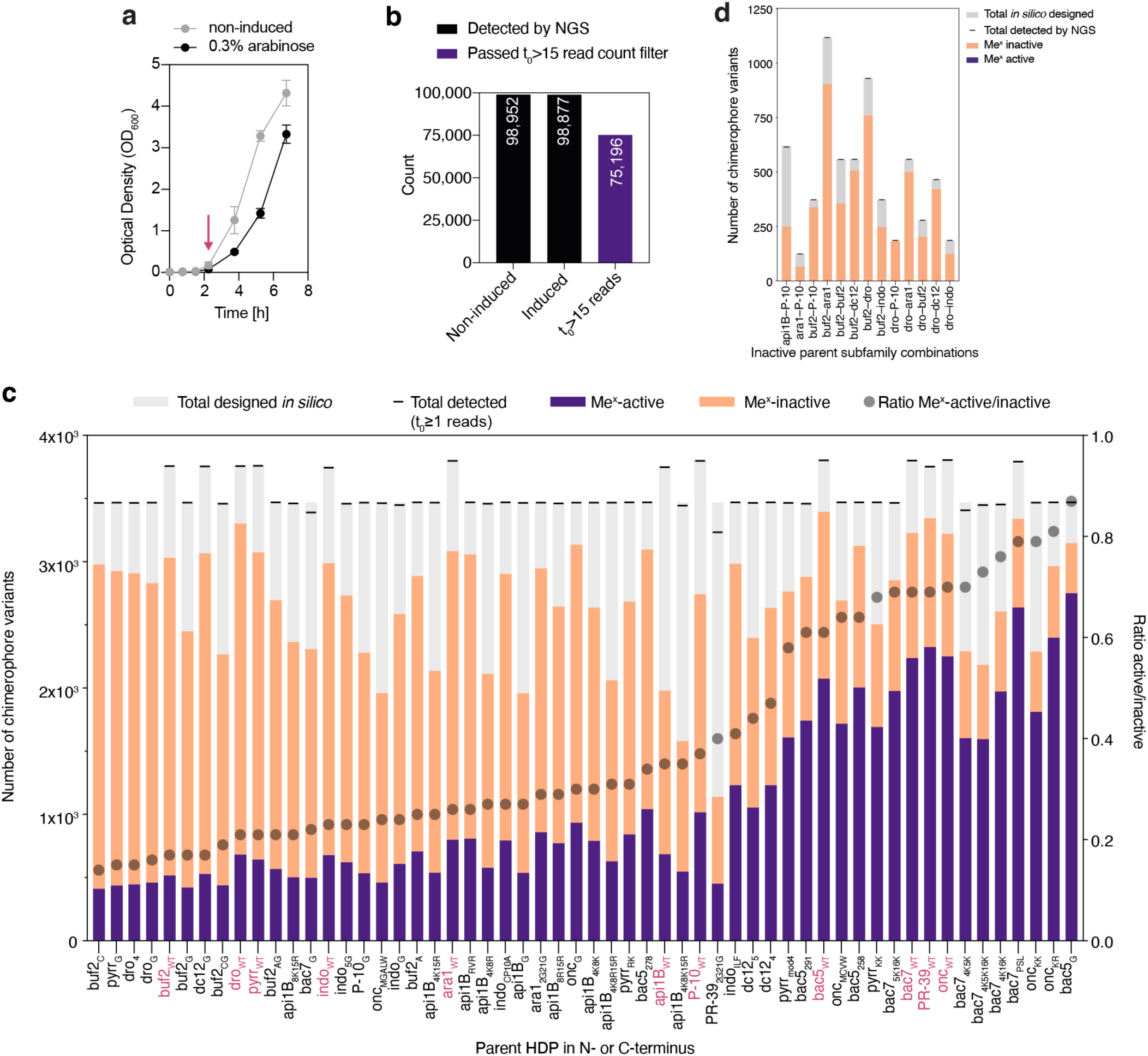
Me^x^ growth curves, library coverage by NGS, and Me^x^-activity ratios. **(a)** Growth curves of *E. coli* TOP10 synthesizing a DNA-encoded chimerophore library, non-induced (gray) and induced (black). Arrow indicates timepoint of peptide synthesis induction with 0.3% (w/v) L-arabinose. **(b)** Total number of chimerophore variants detected by NGS in non-induced samples (gray), induced samples (black), and passing the Me^x^ data quality filter at the point of induction t_0_ > 15 read counts. **(c)** Distribution of NGS-detected (gray), Me^x^-active (purple), and Me^x^-inactive (orange) variants in the library, grouped by parent HDP positioned at either terminus of the chimerophore. Dashed line indicates the total number of chimerophore variants designed *in silico*. Dots represent Me^x^-active-to-inactive ratios per parent. Wild type parents indicated in pink, engineered analogs in black.

**Extended Data Figure 3.**
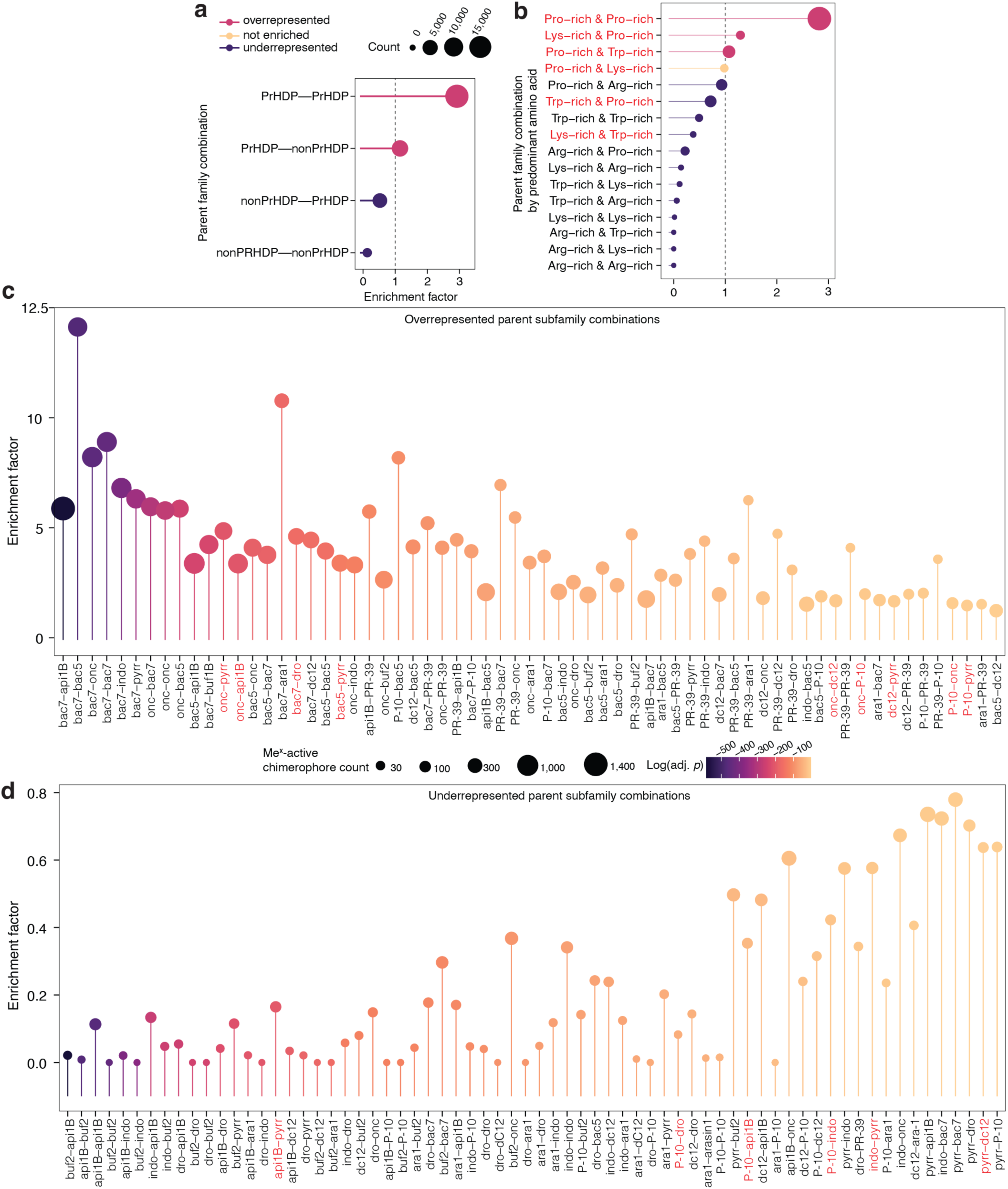
Enrichment analysis of Me^x^-active chimerophores. **(a)** Abundance of parent subfamily combinations in Me^x^-active chimerophores (PrHDP: Pro-rich HDP; non-PrHDP: non-Pro-rich HDPs). **(b)** Abundance of parent family combinations subdivided into predominant amino acid composition groups (PrHDP: Pro-rich; non-PrHDP: disordered Arg-, Trp- and Lys-rich HDPs). **(c)** Significantly overrepresented parent subfamily combinations. Parent subfamilies (e.g., dro) consist of wild-type parent HDPs (e.g., dro_WT_) and their engineered analogs (e.g., dro_4_ and dro_G_). Parent subfamily combinations represented in the 18 selected chimerophores (cp1-cp18) are marked in red. **(d)** Significantly underrepresented parent subfamily combinations. Parent subfamily combinations represented in the 18 selected chimerophores (cp1-cp18) are marked in red.

**Extended Data Figure 4.**
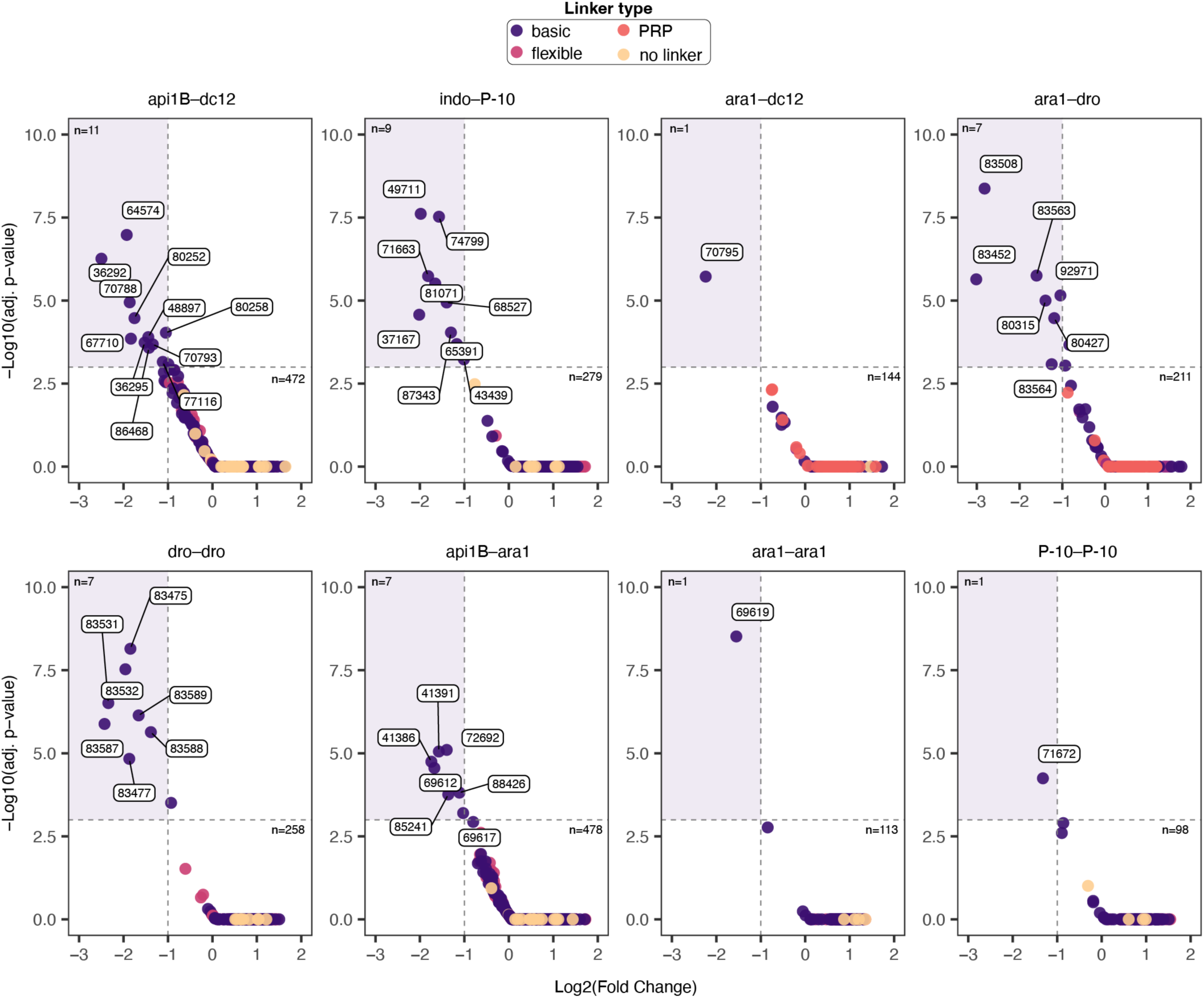
Parent subfamily combinations with activity in Me^x^ exclusively when connected via basic linkers. Dots represent chimerophore variants with basic linkers (purple), flexible linkers (pink), PRP linkers (orange) or without a linker (yellow). Me^x^ activity thresholds are set to LFC < –1, and adjusted p-values < 0.05 (black dashed lines). Quadrants with chimerophore variants that show activity in Me^x^ only with basic linkers are shaded in purple and labelled with their library ID.

**Extended Data Figure 5.**
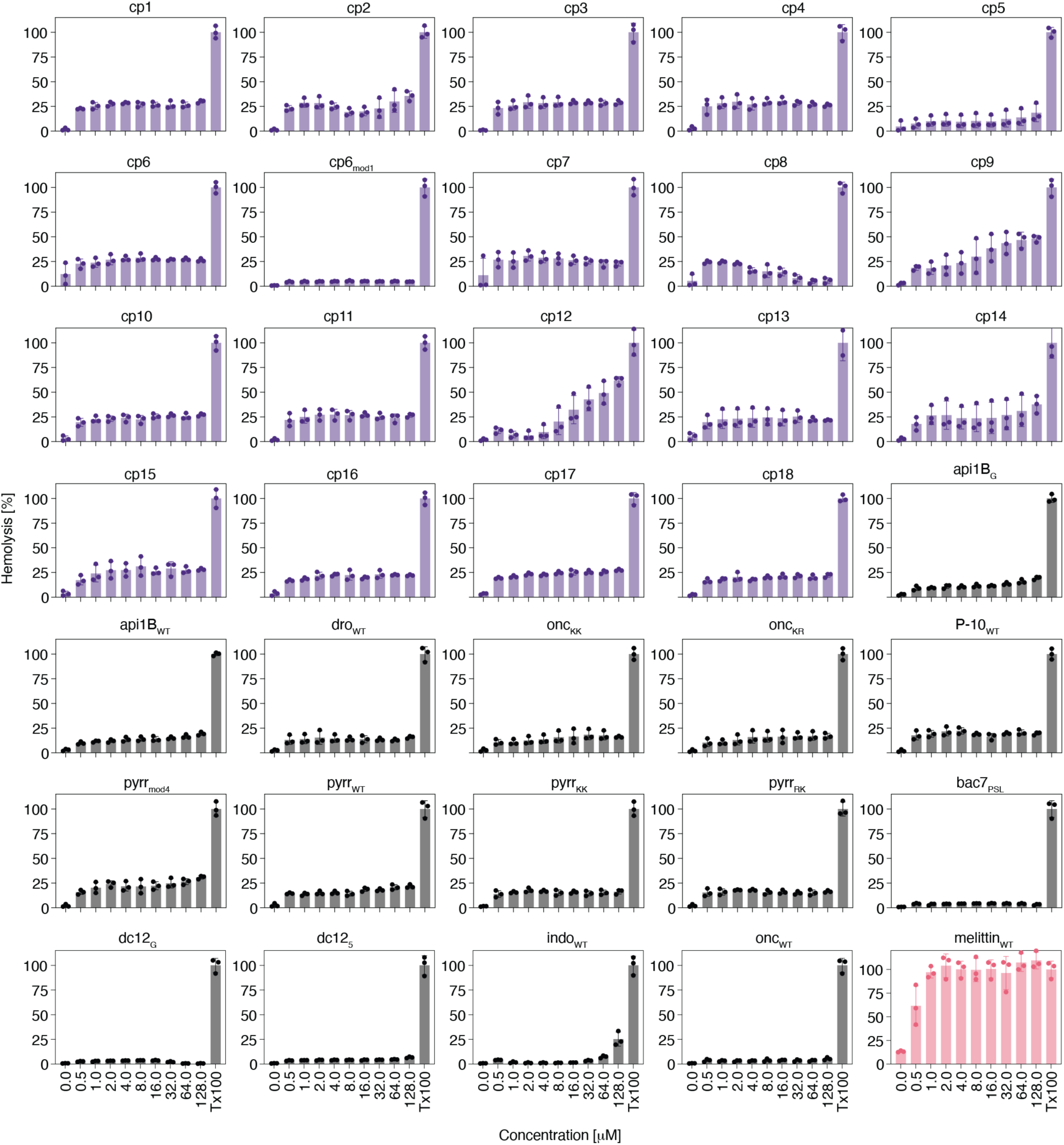
Hemolysis caused by chimerophores and parent HDPs. Hemolysis was measured in microtiter plates via absorbance at 405 nm, in human red blood cells following treatment with 19 chimerophores (cp1-cp18 and cp6_mod1_, purple), 15 parent HDPs (black), and the lytic peptide control melittin (pink). 1% TritonX100 (Tx100) was used as a positive control for complete hemolysis. Bars represent the mean hemolysis relative to positive controls (TX100) and error bars indicate the standard deviation of n=3 biological replicates.

**Extended Data Figure 6.**
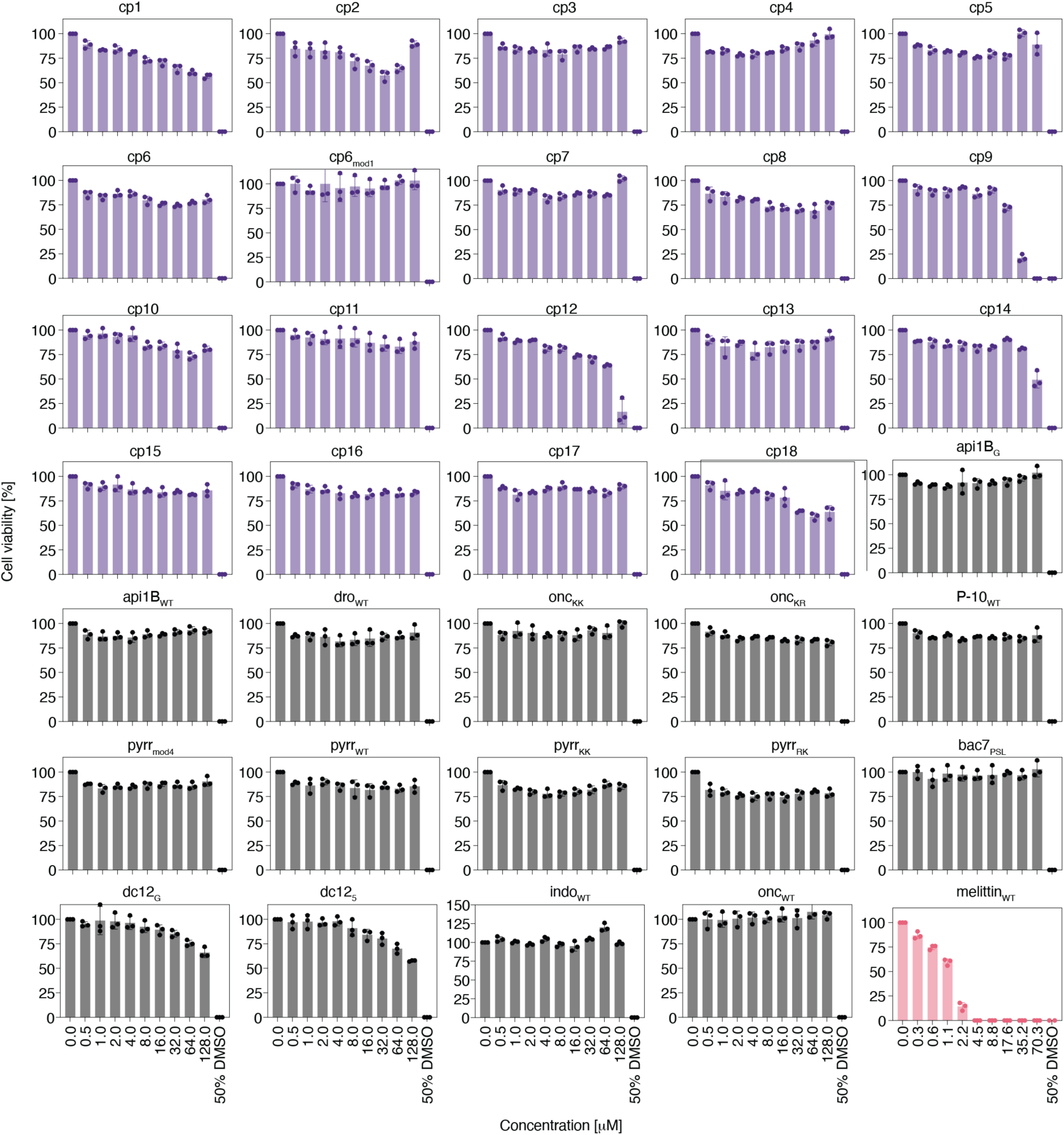
*In vitro* cytotoxicity of chimerophores and parents HDPs. Cell viability was assessed in HEK293 cells using MTT (3-(4,5-dimethylthiazol-2-yl)-2,5-diphenyltetrazolium bromide) assays following treatment with 18 chimerophores (purple), and 15 parents (black), and the lytic peptide control melittin (pink). 50% DMSO was used as a positive control for dead cells. Bars represent the mean cell viability relative to untreated controls (0 µM), with error bars indicating standard deviation of n=3 biological replicates.

**Extended Data Figure 7.**
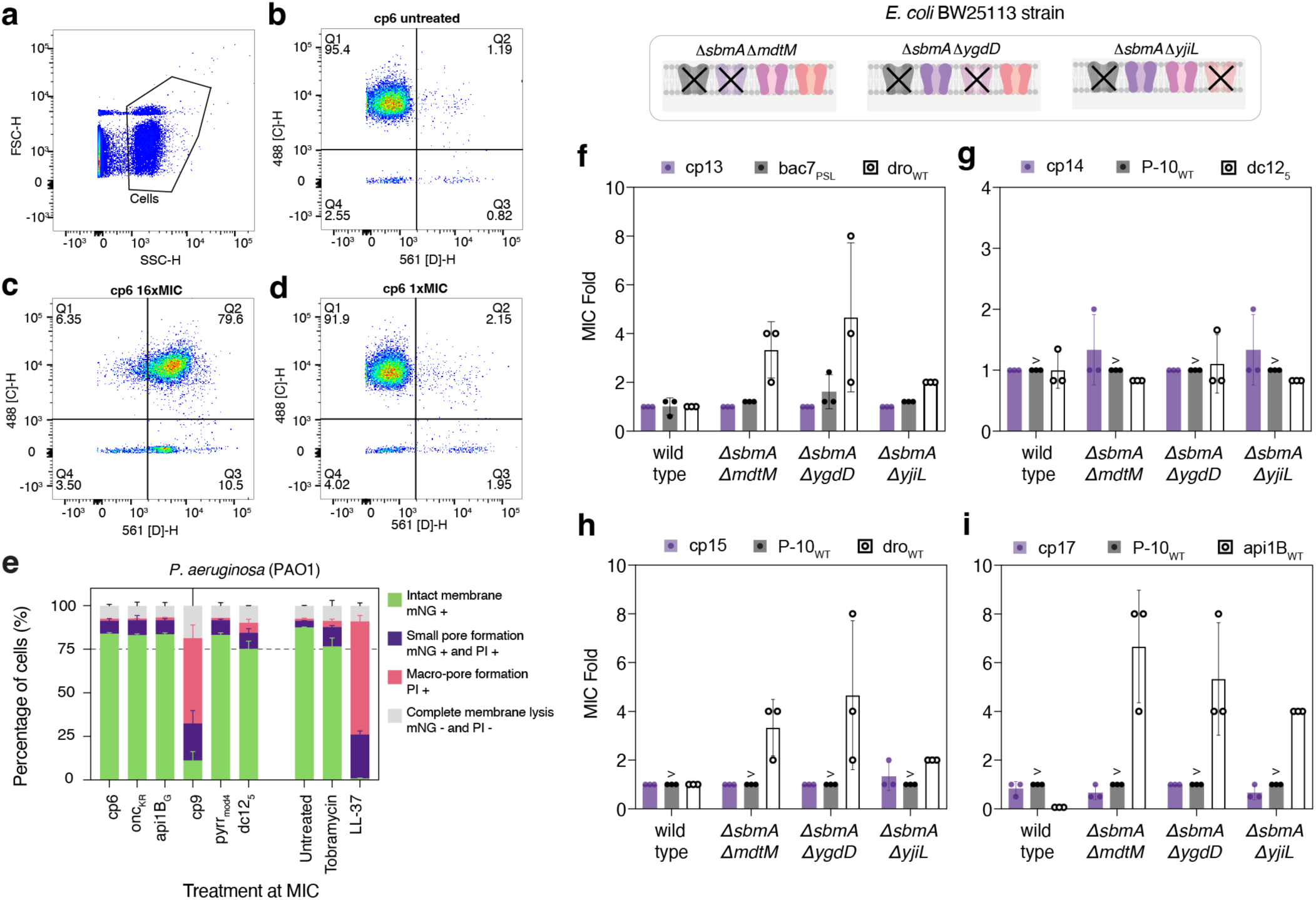
Membrane damage and transport of selected chimerophores in *E. coli* and *P. aeruginosa*. **(a)** Flow cytometry gating strategy for identification of *E. coli* TOP10 cells in membrane damage assays and **(b)** further gates used for categorizing GFP+/PI- (Q1), GFP+/PI+ (Q2), GFP-/PI+ (Q3), and GFP-/PI-(Q4) cells in an untreated control, **(c)** in cp6 treated cells at 16xMIC, **(d)** and at 1xMIC. **(e)** Membrane damage by chimerophores and their parent HDPs (left), and controls (right) including the non-membranolytic antibiotic tobramycin and the membranolytic HDP LL-37 in *P. aeruginosa* PAO1 cells constitutively expressing mNeonGreen. Bars represent the mean of n=2 biological replicates, error bars represent the standard deviation, dashed line indicates lysis threshold of 75% cells. **(f-i)** Activity assays against *E. coli* BW25113 double knockout strains *ΔsbmA ΔmdtM*, *ΔsbmA ΔygdD* and *ΔsbmA ΔyjiL*. Cells were treated with chimerophores (solid fill purple circles) and their parent HDPs, N-terminal parent (black circles, gray bars) and C-terminal parent (open circles). Columns represent the mean and error bars the standard deviation of n=3 biological replicates.

**Extended Data Figure 8.**
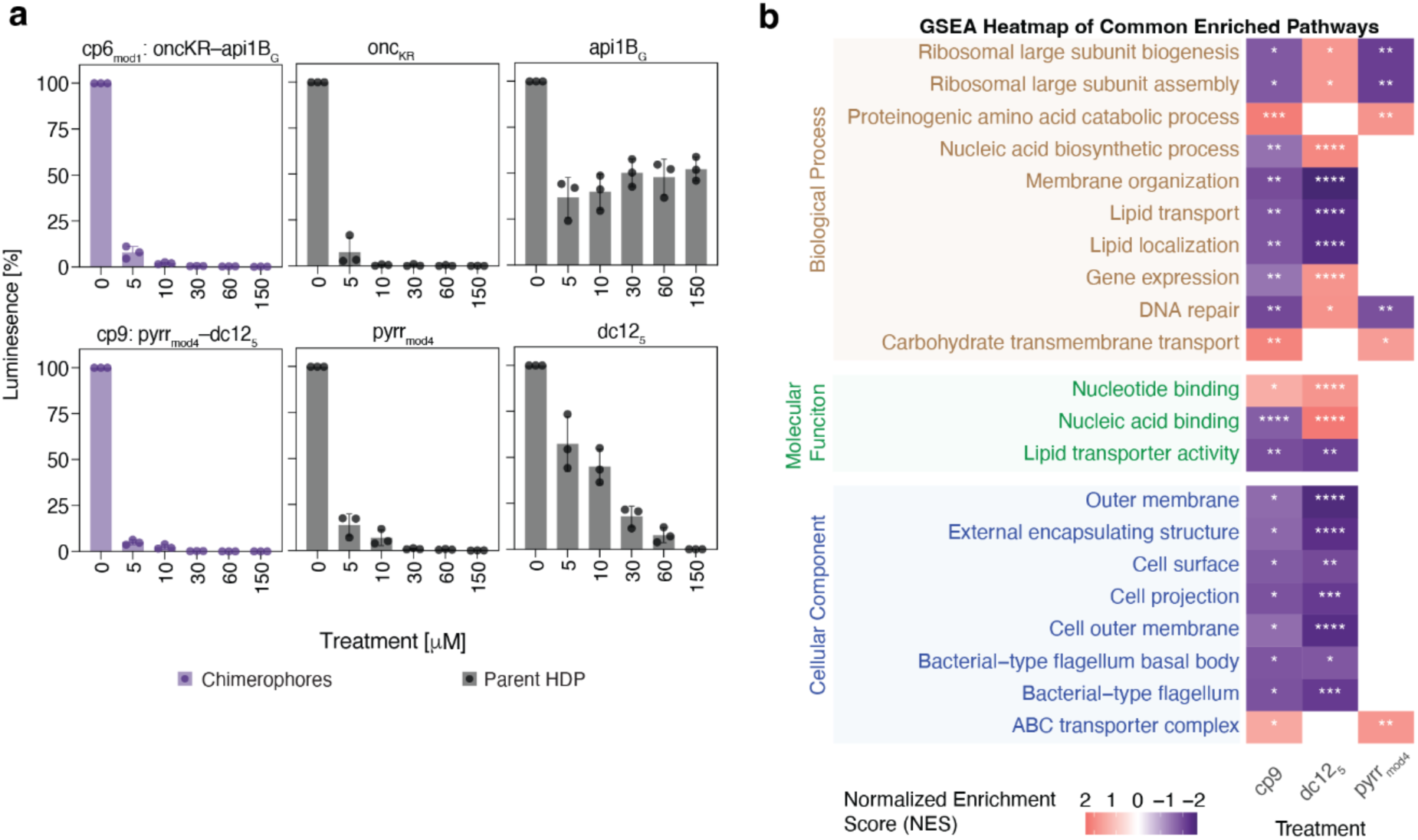
*In vitro* target engagement and interactions of chimerophores and parental HDPs. (a) In vitro transcription translation inhibition assay with increasing concentrations of two lead chimerophores (purple) cp6 and cp9, selected due to high antimicrobial potency and low in vitro toxicity against mammalian cells, and their parent HDPs (black), using luciferase activity as a reporter. The data was normalized to untreated controls (0 µM). Bars represent the mean and error bars represent the standard deviation of the mean of n=3 biological replicates. (b) Gene set enrichment analysis (GSEA) of the *E. coli* ATCC 25922 proteome following PISA treatment with cp9 (16 µg/mL), and its parents, pyrrmod4 (64 µg/mL) and dc125 (16 µg/mL). Heatmap colors indicate the normalized enrichment score (NES), positive values (pink) denote destabilized pathways (reduced solubility/abundance), while negative values (purple) denote stabilized pathways (increased solubility/abundance). Enrichment terms are grouped by gene ontology annotations, biological process (brown), molecular function (green), and cellular component (blue). Statistical significance is indicated by asterisks: * p<0.05, ** p<0.001, *** p<0,0001, **** p<0.00001.

