## Supplementary material for "Chimerophore antibiotics: engineered multimodal host defense peptides"

Suppl. Fig. 1. Selection and phylogenetic analysis of the parent HDP set.

Suppl. Fig. 2. Effect of linker type on $Me^x$-activity of api-bac5 parent subfamily combinations.

Suppl. Fig. 3. AlphaFold predicted 3D models of 18 chimerophores.

Suppl. Fig. 4. Membrane integrity of *E. coli* TOP10 cells treated with chimerophores and their parent HDPs.

Suppl. Fig. 5. Proteome Integral Solubility Alteration (PISA) data quality metrics.

Suppl. Table 1. Parent HDP set information.

Suppl. Table 2. Chimerophore library database **(separate file)**.

Suppl. Table 3. Activity of parent HDPs against *P. aeruginosa* and *S. aureus* in 20% human serum.

Suppl. Table 4. Proteomics data with high/low confidence hits list.

Suppl. Table 5. Genes with mutations identified in whole-genome sequencing of *P. aeruginosa* ATCC 27853 lineages on day 21 of serial passages.

Suppl. Table 6. DNA primers used in this study.

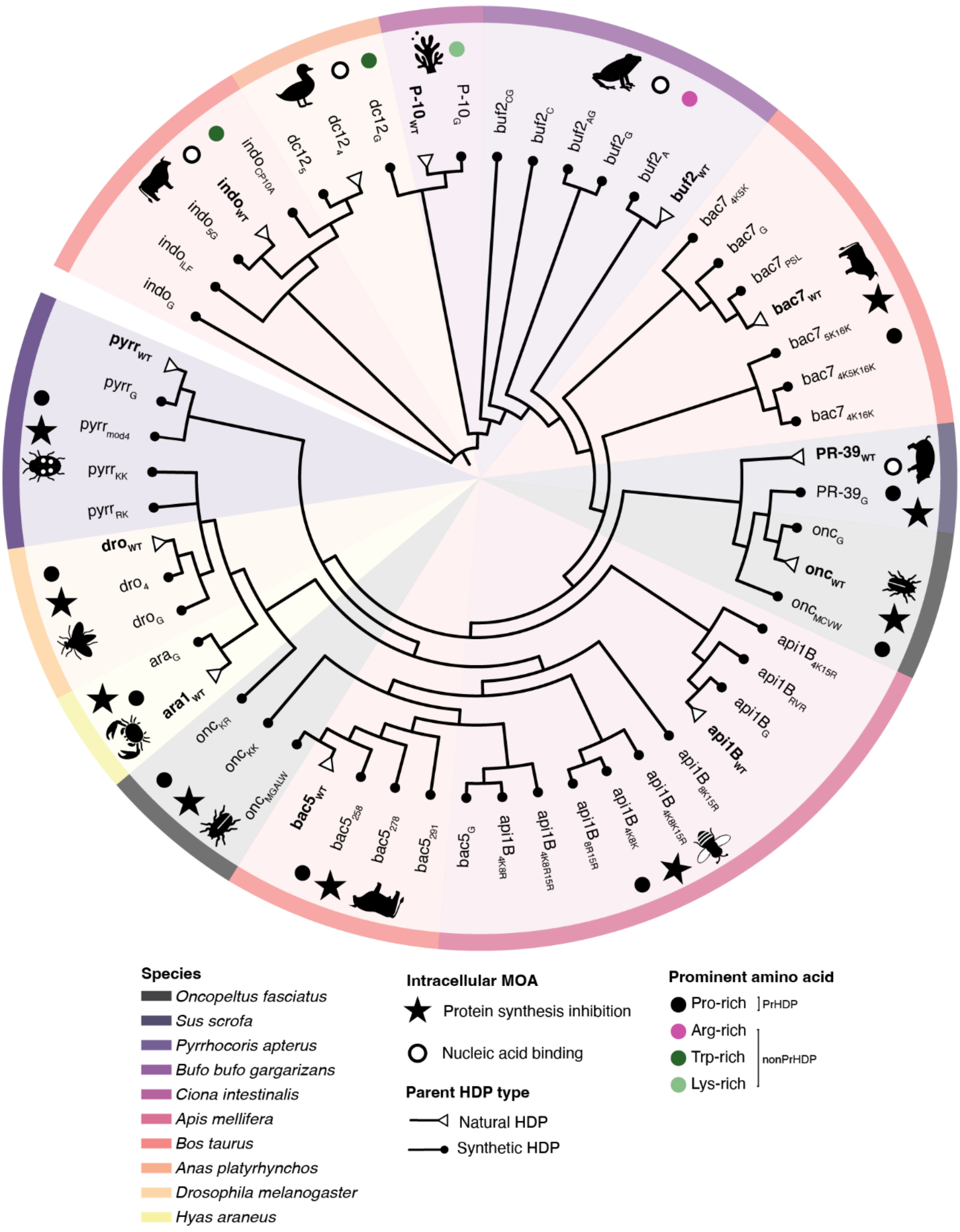


**Supplementary Figure 1. Selection and phylogenetic analysis of the 56 parent HDP set.** Schematic representation of natural (empty triangles) and synthetic (black dots) HDPs selected as the parent set for chimerophore design. The set comprises HDPs derived from 10 diverse species (black icons, background shaded in different colors) categorized by two main intracellular mechanisms of action: protein synthesis inhibition (black stars) and nucleic acid binding (black empty circles). The parent HDPs are also classified into families by prominent amino acid composition: PrHDPs consisting of Pro-rich HDPs (black circles), and non-PrHDPs consisting of Arg-rich (pink circles), Trp-rich (dark green circles), and Lys-rich HDPs (light green circles). Sequences of the 56 HDPs were aligned using MAFFT, curated with BMGE. The tree was inferred using a PhyML model in NGphylogeny.fr, followed by manual trimming and annotation within iTOL.

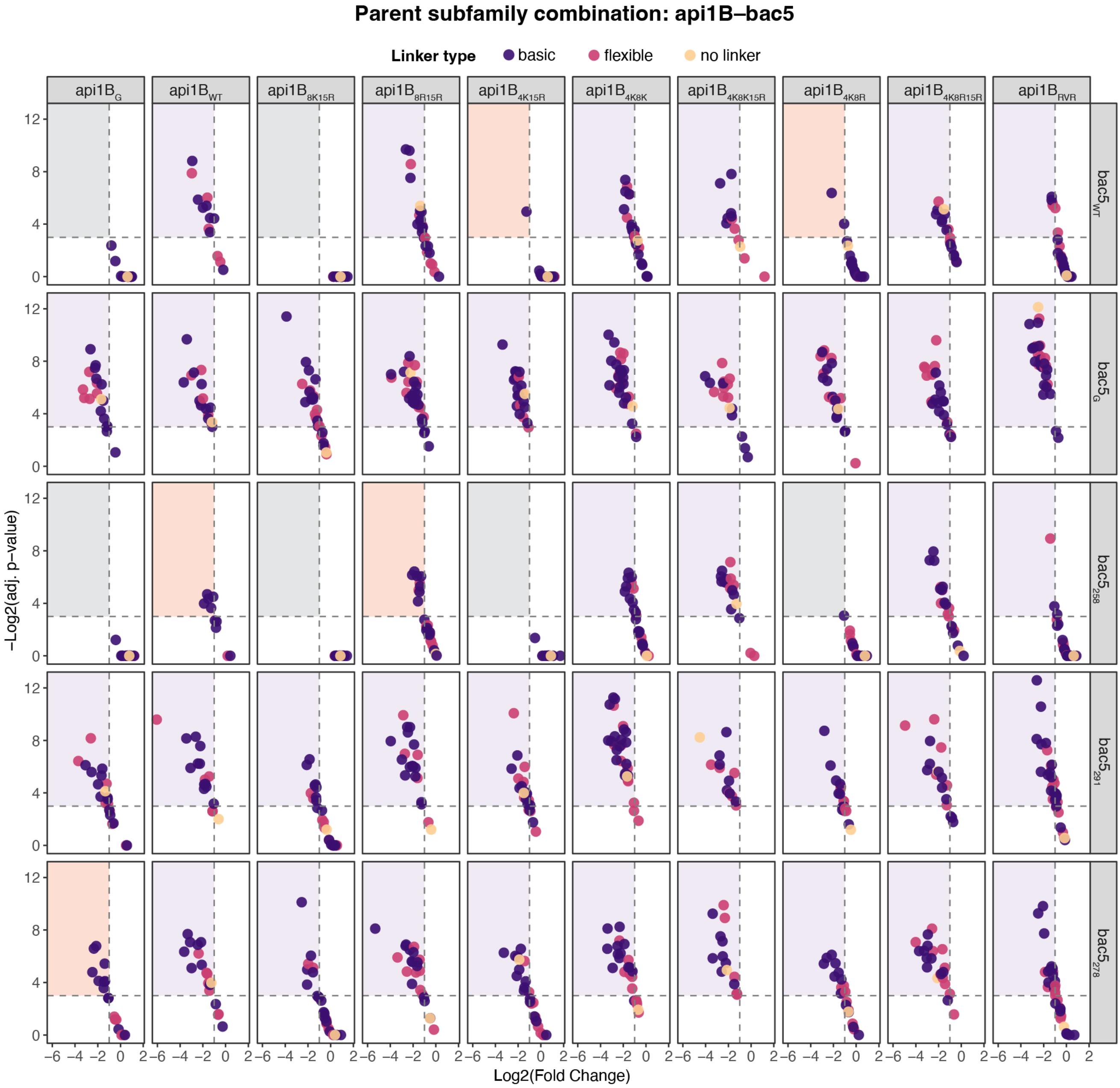


**Supplementary Figure 2. Effect of linker type on Me$^{x}$-activity of api-bac5 parent subfamily combinations.** Parent subfamilies group wild-type parent HDPs (e.g., api1B$_{WT}$) with their respective engineered analogs (e.g., api1B$_{G}$, api1B$_{RVR}$, etc.). The ten api1B-subfamily parents (columns) are shown in combination with five bac5-subfamily parents (rows). Individual dots represent chimerophore variants with different linker types: basic (purple), flexible (pink) or none (yellow). Me$^{x}$-activity thresholds are set to LFC < –1, and adjusted p-values < 0.05 (dashed lines). Quadrants shading denotes Me$^{x}$-activity profiles: orange indicates combinations that are exclusively active with basic linkers, purple indicates activity across multiple linker types, and gray indicates a lack of Me$^{x}$-active variants regardless of linker type.

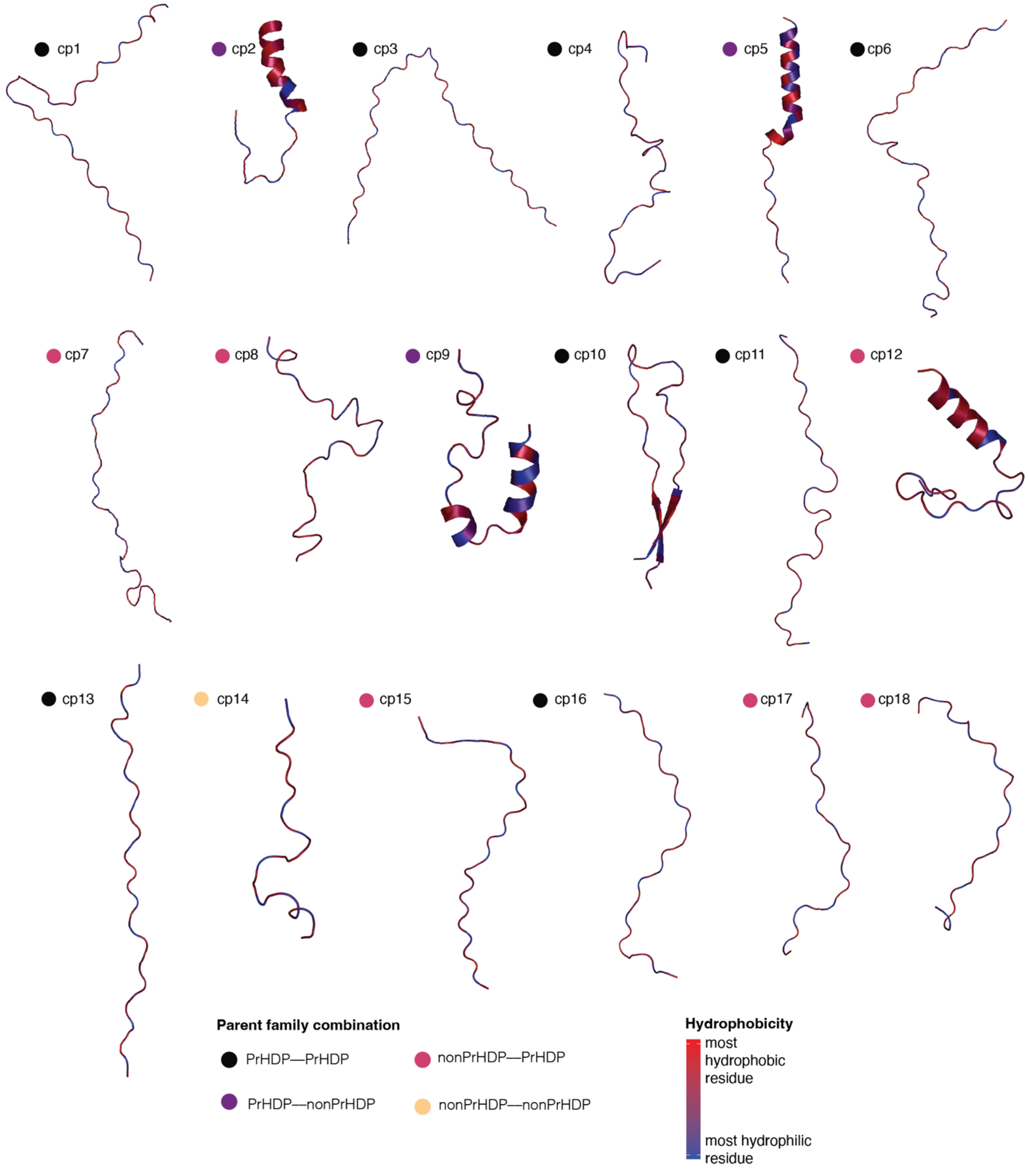


**Supplementary Figure 3. Predicted structural and hydrophobicity profiles of lead chimerophores.** AlphaFold3-predicted 3D models of the 18 chemically synthesized chimerophores, with residues colored according to the Kyte-Doolittle hydrophobicity scale. Blue indicates the least hydrophobic (hydrophilic) residues, while red highlights the most hydrophobic regions. Chimerophores are also classified by parent family combination: dual PrHDP combinations (black circles), PrHDP–non-PrHDP combinations (purple circles), non-PrHDP–PrHDP combinations (pink circles), and dual non-PrHDP combinations (yellow circles). PrHDPs: Pro-rich parents, non-PrHDPs: Trp-rich, Arg-rich or Lys-rich parents.

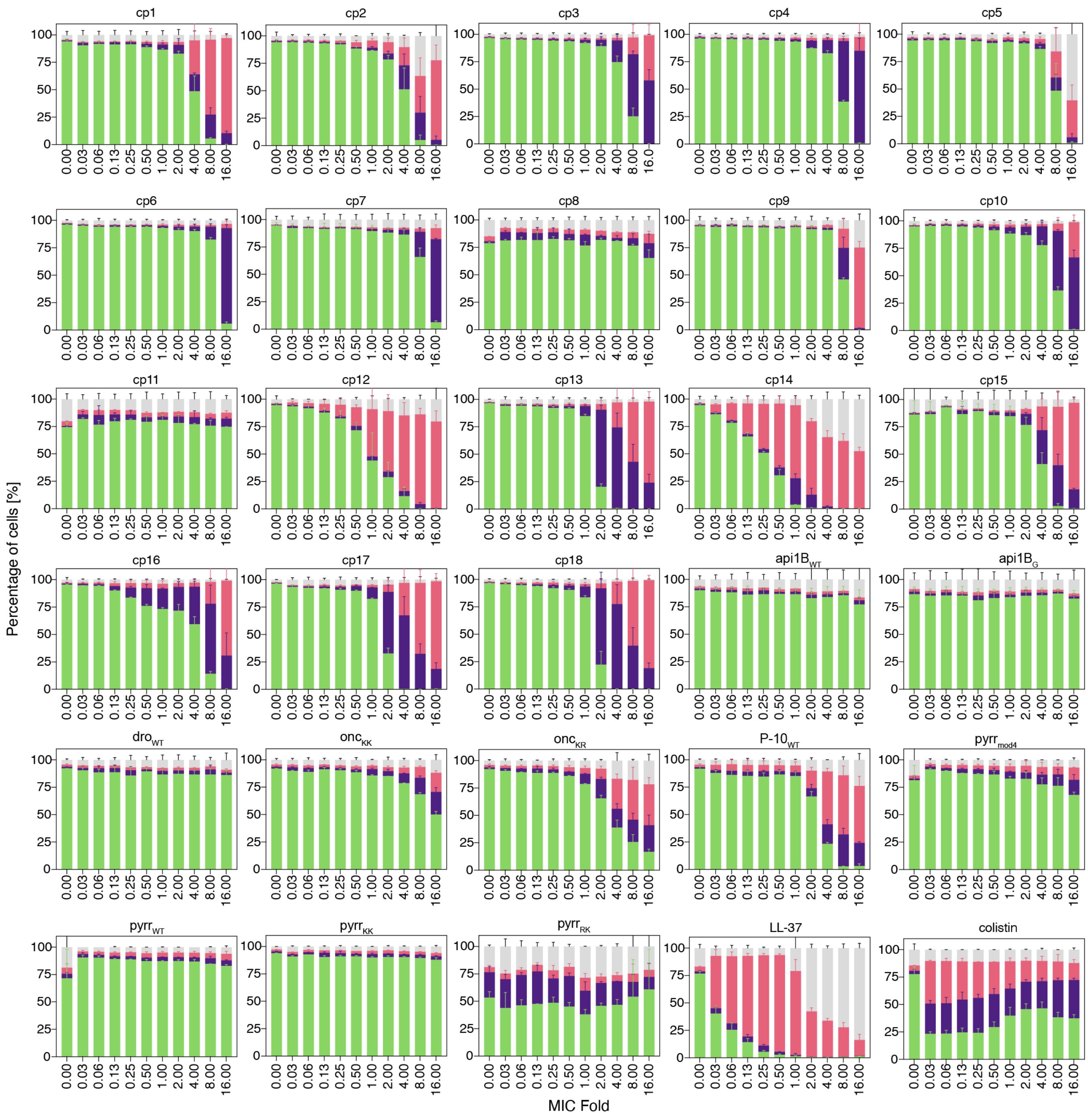


**Supplementary Figure 4. Membrane integrity of *E. coli* TOP10 cells treated with chimerophores and their parent HDPs.** Membrane integrity of *E. coli* TOP10 cells constitutively expressing sfGFP was assessed via flow cytometry, upon treatment of 18 chemically synthesized chimerophores (cp1-cp18), 10 parent HDPs, and the membranolytic HDPs LL-37 and colistin as controls. Peptide treatment was administered at two-fold dilution series starting at 16-fold the MIC; 0.00: untreated controls. Bars represent the mean of n=2 biological replicates, error bars represent the standard deviation.

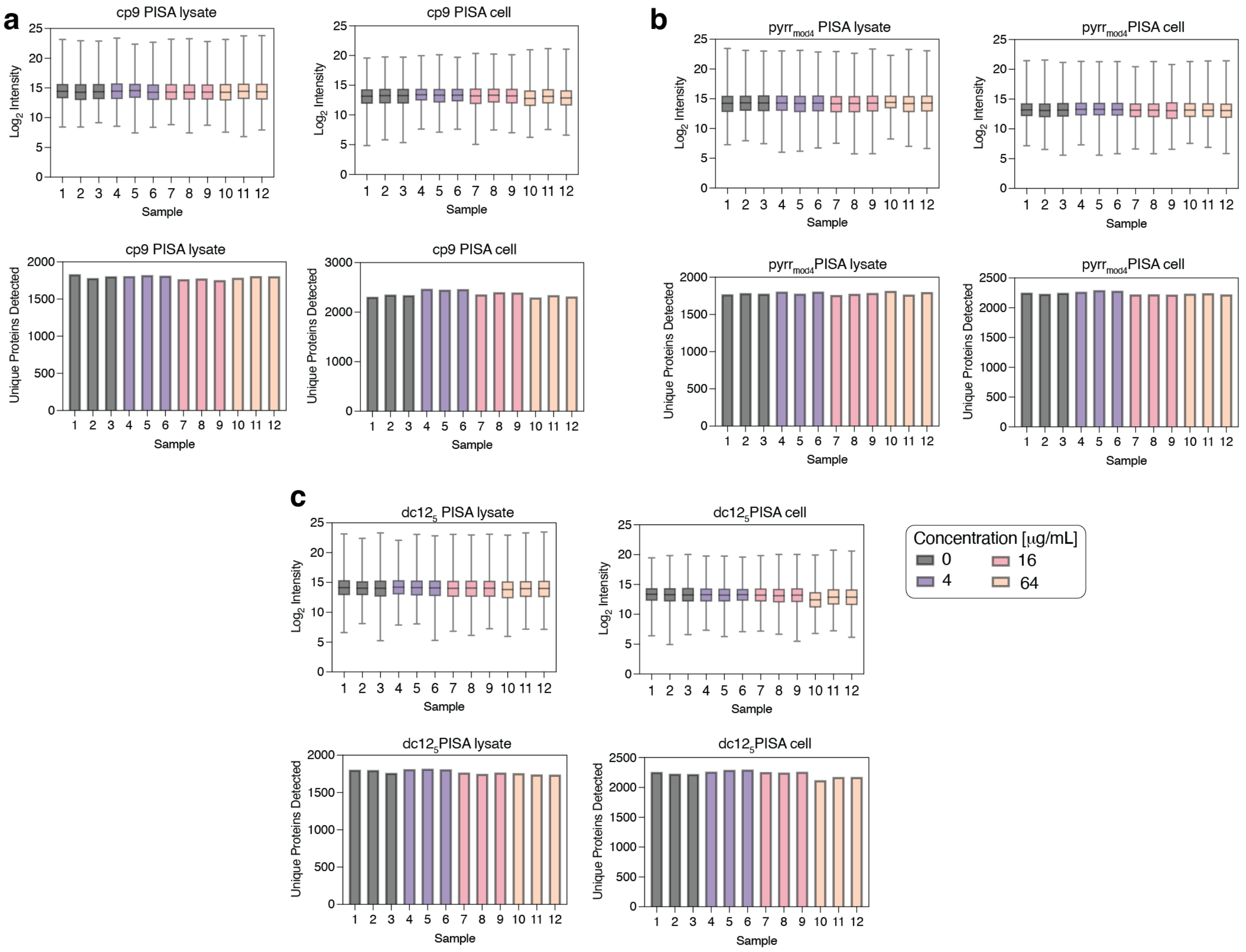


**Supplementary Figure 5. Proteome Integral Solubility Alteration (PISA) data quality metrics.** Global proteomic profiles for *E. coli* ATCC 25922 following treatment with **(a)** cp9, **(b)** pyrr$_{mod4}$ and **(c)** dc12$_5$ at 0 (black), 4 (purple), 16 (pink) and 64 $\mu$g/mL (yellow). Log$_2$ Intensity per sample (top) and unique proteins detected (bottom) are presented for both PISA lysate (left) and whole-cell samples (right). Box plots represent the mean and interquartile range of n=3 biological replicates, error bars indicate the standard deviation of the mean. Bars represent the mean of n=3 biological replicates.

**Supplementary Table 1. Parent HDP set information.** AA: amino acid; MOA: mechanism of action; NB: nucleic-acid binding; TTI: translation termination inhibitor; TEI: translation elongation inhibitor; PSI: protein synthesis inhibitor; TI: type I PrHDP; TII: type II PrHDP. Parent families are indicated by predominant amino acid composition. Amino acid substitutions in engineered analogs are indicated in red.

| *parent subfamily ID* | *parent ID* | *AA sequence* | *length* | *parent family* | *MOA* | *reference* |
|---|---|---|---|---|---|---|
| buf2 | buf2$_{WT}$ | TRSSRAGLQFPVGRVHRLLRK | 21 | Arg-rich | NB | *Park et al. (1996)*[1] |
| buf2 | buf2$_{A}$ | RAGLQFPVGRVHRLLRK | 17 | Arg-rich | NB | *Park et al. (1996)*[1] |
| buf2 | buf2$_{C}$ | RAGLQFPLGRLLRRLLRRLLR | 21 | Arg-rich | NB | *Hao et al. (2013)*[2] |
| buf2 | buf2$_{AG}$ | RAGLQFPVGGVHGLLRK | 17 | Arg-rich | NB | *this work* |
| buf2 | buf2$_{CG}$ | RAGLQFPLGGLLGGLLGGLLR | 21 | Arg-rich | NB | *this work* |
| buf2 | buf2$_{G}$ | TGSSGAGLQFPVGGVHGLLRK | 21 | Arg-rich | NB | *this work* |
| dc12 | dc12$_{4}$ | LIKKIYRKWKRW | 12 | Trp-rich | NB | *Kumar et al. (2020)*[3] |
| dc12 | dc12$_{5}$ | LWKKIYRKWKRW | 12 | Trp-rich | NB | *Kumar et al. (2020)*[3] |
| dc12 | dc12$_{G}$ | LWGGIYGGWGGW | 12 | Trp-rich | NB | *this work* |
| indo | indo$_{WT}$ | ILPWKWPWWPWRR | 13 | Trp-rich | NB | *Selsted et al. (1992)*[4] |
| indo | indo$_{ILF}$ | ILPFKFPFFFPFRR | 13 | Trp-rich | NB | *Subbalakshmi et al.(1996)*[5] |
| Indo | indo$_{CP10A}$ | ILAWKWAWWAWRR | 13 | Trp-rich | NB | *Subbalakshmi et al.(1996)*[5] |
| Indo | indo$_{5G}$ | ILPWGWPWWPWRR | 13 | Trp-rich | NB | *this work* |
| Indo | indo$_{G}$ | ILPWGWPWWPWGG | 13 | Trp-rich | NB | *this work* |
| ara1 | ara1$_{WT}$ | SRWPSPGRPRPFPGRPKPIFRPR | 23 | Pro-rich | PSI | *Stensvåg et al. (2008)*[6] |
| ara1 | ara1$_{2G21G}$ | SGWPSPGRPRPFPGRPKPIFGPR | 23 | Pro-rich | PSI | *this work* |
| bac5 | bac5$_{WT}$ | RFRPPIRRPPIRPPFYP | 17 | Pro-rich TI | TEI | *Mardirossian et al.(2019)*[7] |
| bac5 | bac5$_{258}$ | RFRPPIRRPPIRPPFYR | 17 | Pro-rich TI | TEI | *Mardirossian et al.(2019)*[7] |
| bac5 | bac5$_{278}$ | RWRWPIRRPPIRPPFYR | 17 | Pro-rich TI | TEI | *Mardirossian et al.(2019)*[7] |
| bac5 | bac5$_{291}$ | RWRRPIRRRPIRPPFWR | 17 | Pro-rich TI | TEI | *Mardirossian et al.(2019)*[7] |
| bac5 | bac5$_{G}$ | RFGPPIRGPPIGPPFYP | 17 | Pro-rich TI | TEI | *this work* |
| bac7 | bac7$_{WT}$ | RRIRPRPPRLPRPRPR | 16 | Pro-rich TI | TEI | *Benincasa et al. (2004)*[8] |
| bac7 | bac7$_{5K16K}$ | RRIRKRPPRLPRPRPK | 16 | Pro-rich TI | TEI | *Lai et al. (2019)*[9] |
| bac7 | bac7$_{4K5K}$ | RRIKKRPPRLPRPRPR | 16 | Pro-rich TI | TEI | *Lai et al. (2019)*[9] |
| bac7 | bac7$_{4K16K}$ | RRIKPRPPRLPRPRPK | 16 | Pro-rich TI | TEI | *Lai et al. (2019)*[9] |
| bac7 | bac7$_{4K5K16K}$ | RRIKKRPPRLPRPRPK | 16 | Pro-rich TI | TEI | *Lai et al. (2019)*[9] |
| bac7 | bac7$_{PSL}$ | RRIRPPRLPRPRPRPYFMPRP | 21 | Pro-rich TI | TEI | *Koch et al. (2022)*[10] |
| bac7 | bac7$_{G}$ | RGIRPRPPGLPRPRPR | 16 | Pro-rich TI | TEI | *this work* |

| *parent* | *parent ID* | *AA sequence* | *length* | *parent family* | *MOA* | *reference* |
|---|---|---|---|---|---|---|

| *subfamily ID* | | | | | | |
|---|---|---|---|---|---|---|
| Onc | onc$_{WT}$ | VDKPPYLPRPRPPRRIYNR | 19 | Pro-rich TII | TEI | *Knappe et al. (2010)*[11] |
| Onc | onc$_{MCVW}$ | MDKPCYLPRPRPPVWIYNR | 19 | Pro-rich TI | TEI | *DeJong et al. (2021)*[12] |
| Onc | onc$_{MGALW}$ | MDKPCYGPAPRPPLWIYNR | 19 | Pro-rich TI | TEI | *DeJong et al. (2021)*[12] |
| Onc | onc$_{KK}$ | VDKKPYKPRPRPPPRRIYNR | 19 | Pro-rich TI | TEI | *Lai et al. (2018)*[13] |
| Onc | onc$_{KR}$ | VDKKPYRPRPRPPRRIYNR | 19 | Pro-rich TI | TEI | *Lai et al. (2018)*[13] |
| Onc | onc$_{G}$ | VDGPPYLPRPRPPRGIYNR | 19 | Pro-rich TI | TEI | *this work* |
| PR-39 | PR-39$_{WT}$ | RRRPRPPYLPRPRPPPFFPPRL | 22 | Pro-rich TI | TEI | *Veldhuizen et al. (2014)*[14] |
| PR-39 | PR-39$_{2G21G}$ | RGRPRPPYLPRPRPPPFFPPGL | 22 | Pro-rich TI | TEI | *this work* |
| Pyrr | pyrr$_{WT}$ | VDKGSYLPRPTPPRPIYNRN | 20 | Pro-rich TI | TEI | *Cociancich et al. (1994)*[15] |
| Pyrr | pyrr$_{KK}$ | VDKGSYKPRPTPPKPIYNRN | 20 | Pro-rich TI | TEI | *Lai et al. (2019)*[9] |
| Pyrr | pyrr$_{RK}$ | VDKGSYRPRPTPPKPIYNRN | 20 | Pro-rich TI | TEI | *Lai et al. (2019)*[9] |
| Pyrr | pyrr$_{mod4}$ | VDKGSYLPRPTPTPRYRPIYNRN | 23 | Pro-rich TI | TEI | *Kragol et al. (2002)*[16] |
| Pyrr | pyrr$_{G}$ | VDGGSYLPRPTPPRPIYNGN | 20 | Pro-rich TI | TEI | *this work* |
| api1B | api1B$_{WT}$ | GNNRPVYIPQPRPPHPRL | 18 | Pro-rich TII | TTI | *Casteels et al. (1989)*[17] |
| api1B | api1B$_{RVR}$ | RVRRPVYIPQPRPPHPRL | 18 | Pro-rich TII | TTI | *Taguchi et al. (2009)*[18] |
| api1B | api1B$_{8K15R}$ | GNNRPVYKPQPRPPRPRL | 18 | Pro-rich TII | TTI | *Lai et al. (2019)*[9] |
| api1B | api1B$_{8R15R}$ | GNNRPVYRPQPRPPRPRL | 18 | Pro-rich TII | TTI | *Lai et al. (2019)*[9] |
| api1B | api1B$_{4K15R}$ | GNNKPVYIPQPRPPRPRL | 18 | Pro-rich TII | TTI | *Lai et al. (2019)*[9] |
| api1B | api1B$_{4K8K}$ | GNNKPVYKPQPRPPHPRL | 18 | Pro-rich TII | TTI | *Lai et al. (2019)*[9] |
| api1B | api1B$_{4K8K15R}$ | GNNKPVYKPQPRPPRPRL | 18 | Pro-rich TII | TTI | *Lai et al. (2019)*[9] |
| api1B | api1B$_{4K8R}$ | GNNKPVYRPQPRPPHPRL | 18 | Pro-rich TII | TTI | *Lai et al. (2019)*[9] |
| api1B | api1B$_{4K8R15R}$ | GNNKPVYRPQPRPPRPRL | 18 | Pro-rich TII | TTI | *Lai et al. (2019)*[9] |
| api1B | api1B$_{G}$ | GNNGPVYIPQPRPPHPRL | 18 | Pro-rich TII | TTI | *this work* |
| dro | dro$_{WT}$ | GKPRPYSPRPTSHPRPIRV | 19 | Pro-rich TII | TTI | *Bulet et al. (1993)*[19] |
| dro | dro$_{4}$ | GKPRPYTPRPTSHPRPIRV | 19 | Pro-rich TII | TTI | *Bikker et al. (2016)*[20] |
| dro | dro$_{G}$ | GGPRPYSPRPTSGPRPIGV | 19 | Pro-rich TII | TTI | *this work* |
| P-10 | P-10$_{WT}$ | VSKIKKYLKYKDRI | 14 | Lys-rich | U | *Lu et al. (2014)*[14] |
| P-10 | P-10$_{G}$ | VSGIGGYLGYGDRI | 14 | Gly-rich | U | *this work* |

**Supplementary Table 3. Activity of parent HDPs against *Pseudomonas aeruginosa* and *Staphylococcus aureus* in 20% human serum.**

| | MIC [µg/mL] in 20% human serum | |
|---|---|---|
| **Parent** | *P. aeruginosa* ATCC 27853 | *S. aureus* ATCC 29213 |
| api1$B_G$ | >128 | *NA* |
| api1$B_{WT}$ | >128 | *NA* |
| $dro_{WT}$ | >128 | *NA* |
| $onc_{KK}$ | >128 | *NA* |
| $onc_{KR}$ | >128 | *NA* |
| P-$10_{WT}$ | >128 | *NA* |
| $pyrr_{WT}$ | >128 | *NA* |
| $pyrr_{KK}$ | >128 | *NA* |
| bac$7_{PSL}$ | 9.33 | 64 |
| dc$12_5$ | 5.8 | >128 |
| $onc_{WT}$ | >128 | *NA* |
| $indo_{WT}$ | *NA* | >128 |

*NA: not tested due to inactivity in 25% MHB.*

*Values are the mean of n=6 biological replicates.*

*Only parents with in vitro activity against each strain were tested.*

**Supplementary Table 4.** Proteomics data with high/low confidence hits list. QC: quality control; LFC: $log_2$(fold change).

| *Treatment [16 mg/mL]* | *Protein ID* | *Uniprot Accession Number* | *Gene* | *Description* | *Unique Peptide Count* | *LFC (untreated vs. treated)* | *p-value* | *q-value* | *QC filter passed* | *confidence* |
|---|---|---|---|---|---|---|---|---|---|---|
| **$dc12_5$** | DCM_ECOLI | P0AED9 | dcm | DNA-cytosine methyltransferase | 1 | Inf | NA | 0 | TRUE | low |
| **$dc12_5$** | ELAB_ECOLI | P0AEH5 | elaB | Protein ElaB | 1 | Inf | NA | 0 | TRUE | low |
| **$dc12_5$** | FPG_ECOLI | P05523 | mutM | Formamidopyrimidine-DNA glycosylase | 1 | Inf | NA | 0 | TRUE | low |
| **$dc12_5$** | HEMH_ECOLI | P23871 | hemH | Ferrochelatase | 1 | Inf | NA | 0 | TRUE | low |
| **$dc12_5$** | LEUO_ECOLI | P10151 | leuO | HTH-type transcriptional regulator LeuO | 1 | -Inf | NA | 0 | TRUE | low |
| **$dc12_5$** | MLTC_ECOLI | P0C066 | mltC | Membrane-bound lytic murein transglycosylase C | 1 | Inf | NA | 0 | TRUE | low |
| **$dc12_5$** | SPPA_ECOLI | P08395 | sppA | Protease 4 | 1 | Inf | NA | 0 | TRUE | low |
| **$dc12_5$** | THID_ECOLI | P76422 | thiD | Hydroxymethylpyrimidine/phosphomethylpyrimidine kinase | 1 | Inf | NA | 0 | TRUE | low |
| **$dc12_5$** | UBIC_ECOLI | P26602 | ubiC | Chorismate pyruvate-lyase | 1 | Inf | NA | 0 | TRUE | low |
| **cp9** | ACPH_ECOLI | P21515 | acpH | Acyl carrier protein phosphodiesterase | 1 | Inf | NA | 0 | TRUE | low |
| **cp9** | AROF_ECOLI | P00888 | aroF | Phospho-2-dehydro-3-deoxyheptonate aldolase, Tyr-sensitive | 1 | -Inf | NA | 0 | TRUE | low |
| **cp9** | CORC_ECOLI | P0AE78 | corC | Magnesium and cobalt efflux protein CorC | 15 | 0.92 | 2.30e-04 | 0.046 | TRUE | high |
| **cp9** | CYDA_ECOLI | P0ABJ9 | cydA | Cytochrome bd-I ubiquinol oxidase subunit 1 | 1 | Inf | NA | 0 | TRUE | low |
| **cp9** | DCP_ECOLI | P24171 | dcp | Dipeptidyl carboxypeptidase | 15 | -1.71 | 8.91e-07 | 0.002 | TRUE | high |
| **cp9** | GAL1_ECOLI | P0A6T3 | galK | Galactokinase | 17 | -0.80 | 1.72e-04 | 0.046 | TRUE | high |
| **cp9** | HFQ_ECOLI | P0A6X3 | hfq | RNA-binding protein Hfq | 10 | 3.050 | 2.96e-06 | 0.003 | TRUE | high |
| **cp9** | HHA_ECOLI | P0ACE3 | hha | Hemolysin expression-modulating protein Hha | 3 | -Inf | NA | 0 | TRUE | low |
| **cp9** | NADE_ECOLI | P18843 | nadE | NH(3)-dependent NAD(+) synthetase | 16 | -0.62 | 2.25e-04 | 0.046 | TRUE | high |
| **cp9** | PRMB_ECOLI | P39199 | prmB | 50S ribosomal protein L3 glutamine methyltransferase | 6 | -0.89 | 2.4905e-04 | 0.046 | TRUE | high |

| | | | | | | | | | | |
|---|---|---|---|---|---|---|---|---|---|---|
| **cp9** | RLMD_ECOLI | P55135 | rlmD | 23S rRNA (uracil(1939)-C(5))-methyltransferase RlmD | 1 | Inf | NA | 0 | TRUE | low |
| **cp9** | RL32_ECOLI | P0A7N4 | rpmF | 50S ribosomal protein L32 | 5 | -1.15 | 8.73e-05 | 0.041 | TRUE | high |
| **cp9** | RS21_ECOLI | P68679 | rpsU | 30S ribosomal protein S21 | 8 | 1.63 | 8.14e-05 | 0.041 | TRUE | high |
| **cp9** | RUVC_ECOLI | P0A814 | ruvC | Crossover junction endodeoxyribonuclease RuvC | 4 | -0.76 | 1.76e-04 | 0.046 | TRUE | high |
| **cp9** | YADG_ECOLI | P36879 | yadG | Uncharacterized ABC transporter ATP-binding protein YadG | 4 | 1.66 | 1.30e-04 | 0.046 | TRUE | high |
| **cp9** | YBJP_ECOLI | P75818 | ybjP | Uncharacterized lipoprotein YbjP | 2 | Inf | NA | 0 | TRUE | low |
| **cp9** | YDCY_ECOLI | P64455 | ydcY | Uncharacterized protein YdcY | 1 | Inf | NA | 0 | TRUE | low |
| **cp9** | YNCD_ECOLI | P76115 | yncD | Probable TonB-dependent receptor YncD | 2 | Inf | NA | 0 | TRUE | low |
| **cp9** | YOAA_ECOLI | P76257 | yoaA | Probable ATP-dependent DNA helicase YoaA | 1 | Inf | NA | 0 | TRUE | low |
| **$pyrr_{mod4}$** | LPTB_ECOLI | P0A9V1 | lptB | Lipopolysaccharide export system ATP-binding protein LptB | 1 | Inf | NA | 0 | TRUE | low |
| **$pyrr_{mod4}$** | MIAA_ECOLI | P16384 | miaA | tRNA dimethylallyltransferase | 1 | -Inf | NA | 0 | TRUE | low |
| **$pyrr_{mod4}$** | MLTB_ECOLI | P41052 | mltB | Membrane-bound lytic murein transglycosylase B | 1 | Inf | NA | 0 | TRUE | low |
| **$pyrr_{mod4}$** | NFI_ECOLI | P68739 | nfi | Endonuclease V | 1 | -Inf | NA | 0 | TRUE | low |

**Supplementary Table 5.** Genes that displayed mutations in whole-genome sequencing of most resistant *P. aeruginosa* lineages from serial passage day 21. The NCBI locus tag is based on the reference genome of *P. aeruginosa* PAO1 (GenBank: GCA_000006765.1). Only the genes with reported function are referenced. Unreferenced genes are predicted based on sequence homology to characterized gene products. *: premature stop codon; Ext*: extended stop codon; fs*: frameshift; D: deletion.

| *Treatment* | *Gene* | *Locus tag (PAO1)* | *Gene product* | *Involved in* | Mutations identified |
|---|---|---|---|---|---|
| meropenem | *oprD* | PA0958 | Outer membrane porin D | Transmembrane transport; carbapenem resistance[21] | Trp277* |
| | *nalD* | PA3574 | TetR-family transcriptional repressor | Controls the MexAB-OprM multidrug efflux pump[22] | Thr43Pro |
| | *gyrA* | PA3168 | DNA gyrase subunit A | DNA-templated DNA replication; Fluoroquinolone resistance[23] | Leu591His |
| | *yedA* | PA4783 | Putative drug/metabolite pump | Unknown function | Val74Gly |
| | *pyrG* | PA3637 | CTP synthetase | Pyrimidine nucleotide biosynthetic process | Lys541Gluext* |
| Parent mixtures: | | | | | |
| $onc_{KR}$ + $api1B_{G}$ | *yedA* | PA4783 | Putative drug/metabolite pump | Unknown function | Val74Gly |
| $pyrr_{mod4}$ + $dc12_{5}$ | *yedA* | PA4783 | Putative drug/metabolite pump | Unknown function | Val74Gly |
| Chimerophores: | | | | | |
| cp6 | *PA1810* | PA1810 | ATP-binding cassette (ABC) transporter; Solute-binding protein family 5 domain-containing protein | Oligopeptide transport; transmembrane transport | Trp237* |
| | *ispC (dxr)* | PA3650 | 1-deoxy-D-xylulose 5-phosphate reductoisomerase | Isoprenoid biosynthetic process; antibiotic resistance in *A. baumannii*[24] | Asp360Gly |
| | *desT* | PA4890 | Transcriptional regulator | Negative regulation of DNA-templated transcription and fatty acid metabolic process[25] | Thr74Pro |
| | *yedA* | PA4783 | Putative drug/metabolite pump | Unknown function | Val74Gly |
| cp9 | *PA1288* | PA1288 | Hypothetical outer membrane protein transport protein | Long-chain fatty acid transport[26] | Tyr143Ilefs*1 |
| | *pslD* | PA2234 | Biofilm formation protein PslD | Polysaccharide biosynthesis/export; biofilm formation[27] | ΔMet-Leu-Ala10 |
| | *yedA* | PA4783 | Putative drug/metabolite pump | Unknown function | Val74Gly |

### Supplementary Table 6. DNA primers used in this study.

| *Primer name* | *Sequence (5' to 3')* | *Description* |
|---|---|---|
| Primer 1 | TGTCTGCAGAGGAGATATAAATG | $Me^x$ forward primer[28] |
| Primer 2 | GCCCTGCACAAAGCTTAC | $Me^x$ reverse primer[28] |
| mdtM-fw | GATTAACCGCAAATTTTAATTACAC | Knockout confirmation |
| mdtM-rev | CAATCCGGTCTTTATTACGC | Knockout confirmation |
| ygdD-fw | GAGTGGCGCTGGCAAATAAC | Knockout confirmation |
| ygdD-rev | GTAATACGATCTTCTGGCGG | Knockout confirmation |
| yjiL-fw | CTGAAAATGCTCAGCCAGATG | Knockout confirmation |
| yjiL-rev | GAATCACAATAATGGAGATGC | Knockout confirmation |
